# Deep behavioral phenotyping of pathogen infected mosquitoes reveals species-specific behavior changes enhancing transmission

**DOI:** 10.64898/2026.01.06.697763

**Authors:** Zhong Wan, Theo Maire, Emilie Giraud, Vladyslav Kalyuzhnyy, Geert-Jan van Gemert, Kjerstin Lanke, Rianne Stoter, Benjamin Mordmüller, Teun Bousema, Louis Lambrechts, Felix J.H. Hol

**Author notes:** these authors contributed equally to this work (Theo Maire, Emilie Giraud). Institut Pasteur, Université Paris Cité, Chemogenomic and Biological Screening Core Facility, C2RT, CNRS UMR3523, Paris, France.

## Abstract

Mosquito-borne pathogens are often proposed to alter mosquito behavior to enhance transmission, yet the prevalence, magnitude, and consequences of such effects remain unclear. Using custom high-throughput behavioral assays and deep-learning analysis, we quantified the blood-feeding behavior of more than 5,000 mosquitoes, and long-term activity patterns of more than 1,000 mosquitoes across six clinically relevant mosquito–pathogen combinations. Infection effects were highly species-specific. *Anopheles stephensi* infected with an African *Plasmodium falciparum* strain showed elevated flight activity throughout infection, an increased tendency to blood-feed upon host contact, and prolonged feeding times. In contrast, infection with an Asian *P. falciparum* strain or *P. vivax* had little impact on *An. stephensi* behavior. We detected no behavioral changes associated with *P. falciparum* infection in *Anopheles gambiae s.s.* or *Anopheles coluzzii.* Interestingly, dengue virus infection in *Aedes aegypti* increased landing, probing, and engorgement rates. Together, these results demonstrate that infection-induced behavioral change is not a general phenomenon among mosquito-borne pathogens, yet differs markedly per mosquito and pathogen species. A transmission model parameterized with our data showed that even modest behavioral shifts can substantially increase transmission, underscoring the potential epidemiological importance of subtle phenotypic shifts.

## Main

In pursuit of a blood meal, mosquitoes put approximately half the world population at risk for some of the most relevant human pathogens including malaria parasites and dengue virus, cumulatively resulting in approximately 600,000 deaths annually^1^. The tremendous impact mosquitoes have on society is rooted in their unique behavioral characteristics, that make them exceptionally effective at transmitting pathogens. Blood feeding, in which pathogens may be transferred to a human via the mosquito’s saliva, is a crucial step in transmission. Detailed knowledge of this behavior is therefore essential to understand the transmission dynamics of mosquito-borne pathogens, and necessary to design effective control strategies.

Successful blood feeding is contingent on navigating a variety of host-associated sensory cues including carbon dioxide, heat, and body odor ^2–5^. The sensory integration of these cues enables mosquitoes to perform highly efficient and selective host-seeking behavior, and depends on the physiological status of the mosquito^6–8^. Pathogen infection changes mosquito physiology fundamentally, including metabolic, neurological, and immunological processes^9–12^. Given the intimate link between physiology and behavior, these pathogen-induced changes may translate into altered behaviors ultimately affecting pathogen transmission.

Pathogen-induced changes in mosquito behavior have indeed been hypothesized to contribute to the efficiency of mosquito-borne pathogen transmission. Field collections of indoor resting mosquitoes gave credence to this hypothesis as *P. falciparum* infection in certain *Anopheles* species was associated with an increase in multiple feeding, possibly enhancing transmission through increased mosquito-human contact^13,14^. Experimental studies suggested enhanced attraction to human odor^15,16^ in *Plasmodium* infected *Anopheles* mosquitoes, and reported increased probing^17^ and biting persistence^18,19^. The majority of these studies, however, used rodent-specific *Plasmodium* species, challenging interpretations towards the human context. Indeed, rodent-specific parasites have different effects on mosquito responses to odors compared to human-specific parasites, casting doubt on the relevance of previous work with rodent-specific parasites^20^. Moreover, reports of parasite-induced behavioral changes are contrasted by several studies reporting no such changes in *P. falciparum* infected *Anopheles*^21,22^. Given these discrepancies, the nature, magnitude, and species specificity of *Plasmodium*-induced behavioral changes in *Anopheles* mosquitoes remains an open question. This situation is mirrored by arbovirus infections in *Aedes* and *Culex* mosquitoes. The impact of arboviruses on mosquito behavior has been studied across a range of mosquito-virus combinations^23–29^, yet behavioral changes are often subtle, differ between virus and mosquito species, and agreement among studies is limited ^30^. Extracting a coherent view from the available literature is not only challenging because of the variety of mosquito and pathogen species used (including non-human pathogens), but is further complicated by the plethora of different methods and approaches used to characterize behaviors, preventing direct comparisons of studies or combined analyses.

The rapid and intricate movements involved in mosquito blood feeding pose significant challenges to the quantitative assessment of blood feeding and biting dynamics. Probing, the process where a mosquito briefly inserts its minute mouthparts into the skin to search for a blood vessel, is challenging to quantify yet highly relevant for pathogen transmission even in the absence of blood engorgement^31,32^. Furthermore, conventional methods to study mosquito biting and engorging behaviors depend upon direct host contact, requiring laboratory animals, limiting throughput and sample size, and preventing detailed automated behavioral characterization. To overcome these limitations, we recently developed innovative approaches enabling automated assessment of long-term trends in mosquito activity patterns, and detailed investigations of mosquito biting and engorging dynamics^33–35^. Advances in deep-learning based tools for the automatic quantification of animal behavior^36–38^, combined with experimental arenas designed to study particular behaviors have been crucial in this development.

Here, we leverage the advantages of high-resolution video tracking and deep-learning based behavioral quantification, to characterize short-range host-seeking, biting, and engorging behavior, and long-term activity patterns and sugar feeding behavior across two major human pathogens and six clinically-relevant mosquito-pathogen combinations. We collected population- and individual-level behavioral statistics derived from over 5000 mosquitoes to gain a robust understanding of the behavioral impact of different human pathogens on their mosquito vectors. Our results reveal that some pathogens have subtle, yet important effects on the behavior of their mosquito host, while other mosquito-pathogen combinations show no measurable behavioral effects of infection. These results suggest that pathogen-induced behavioral manipulation may be much less pervasive among mosquito-borne pathogens than previously thought. Mathematical modeling shows that in the systems where we do observe changes, the observed behavioral alterations may have a significant impact on pathogen transmission, despite the modest magnitude of behavioral change.

## Results

### Shifts in basal activity patterns are rare among mosquito-pathogen combinations

After being taken up in a blood meal, pathogens commence on a journey infecting the midgut, disseminating to the hemocoel, to eventually reach the salivary glands. While *Plasmodium* parasites and dengue virus (DENV) have a distinctly different infection biology in the mosquito, under a standard rearing conditions, both pathogens complete the journey from the midgut to the salivary glands in approximately 10-14 days^39–41^. During this extrinsic incubation period (EIP), pathogen load increases dramatically and marked shifts occur in mosquito physiology. We asked if this dynamic process has an effect on mosquito activity patterns and sugar feeding behavior. From the pathogen’s perspective, it has been hypothesized that flight activity may be reduced during the non-infectious stage (pre-salivary gland), whereas a pathogen could benefit from increased activity when it has reached the salivary glands. To test this hypothesis we deployed BuzzWatch, a novel experimental platform designed to characterize long-term activity patterns of free flying mosquitoes in a light cycle controlled environment (**Fig. 1**).^35^ We quantified behavioral parameters, including flight activity patterns, the duration and speed of flight bouts, and sugar feeding activity, of cohorts of 40 mosquitoes continuously from the time mosquitoes received the infectious blood meal, until 21 days post infection (dpi). Control mosquitoes received a non-infectious blood meal.

**Figure 1.**
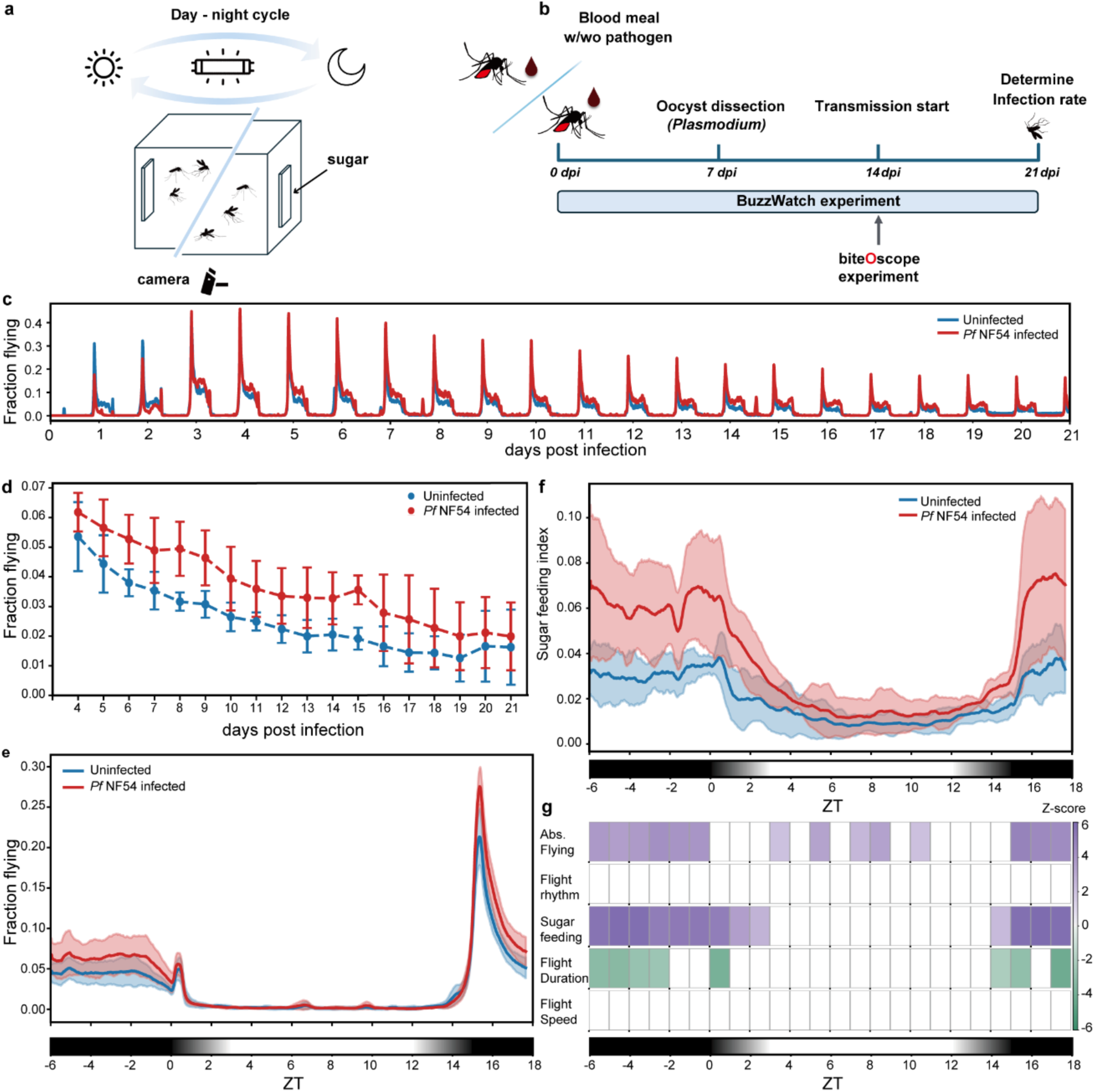
Long-term monitoring of mosquito flying and sugar feeding behavior. **a.** Schematic of BuzzWatch assay. Forty female mosquitoes receiving a pathogen-infected or control blood meal are housed in a transparent cage with continuous access to sugar water. A camera mounted underneath the cage records mosquito behavior continuously. **b.** Timeline of mosquito infections and experiments. Mosquitoes receive a pathogen infected or uninfected control blood meal at 0 dpi. BuzzWatch experiments commence at 0 dpi and are terminated at 21 dpi. To assess prevalence and intensity of *Plasmodium* infections, a subset of mosquitoes is used for midgut dissection to determine oocyst prevalence and number at 7 dpi. BiteOscope experiments are performed at 14 dpi when both *Plasmodium* and DENV reach peak infectiousness. Mosquitoes used for biteOscope experiments are sacrificed immediately upon terminating the experiment at 14 dpi, while mosquitoes used in BuzzWatch experiments were sacrificed at 21 dpi. Upon conclusion of both experiments, infection prevalence of each cohort was determined **c.** Instantaneous fraction of flying mosquitoes over the 3-week period recorded by in the BuzzWatch assay comparing *An. stephensi* infected with *Pf* NF54 (red) and uninfected controls (blue) (mean of *n* = 3 replicates). Curves show the typical nocturnal activity pattern of *Anopheles* mosquitoes. **d.** Mean flight activity per day (*n* = 3) reveals that increased flight activity in *Pf* NF54 infected *An. stephensi* persists for multiple weeks post infection. Days 0-3 post blood meal are excluded as activity is low during this period due to blood digestion. **e.** Averaging activity curves of all 21 days (*n* = 3 replicates) shows that the shape of the activity pattern is similar between *Pf* NF54 infected *An. stephensi* and uninfected controls, yet infected *An. stephensi* exhibit increased flight activity compared to controls across the entire active period. **f.** Daily average of the sugar feeding index over the 3-week experimental period shows significantly increased nocturnal sugar feeding for *Pf* NF54 infected *An. stephensi*. **g.** Heatmap of GLMM test for 6 different quantitative metrics characterizing mosquito flight and sugar feeding behavior. Color scale reflects z-scores (purple = positive, green = negative), only z-scores with statistical significance (p < 0.05) are displayed.

Regardless of infection status, all mosquito species display their typical activity pattern, with nocturnal activity for *Anopheles* mosquitoes, and diurnal activity for *Ae. aegypti*. While in most of the mosquito-pathogen combinations, activity patterns did not differ between infected and non-infected mosquitoes, *An. stephensi* infected with *P. falciparum* strain NF54 (*Pf* NF54, African origin) exhibited a significantly increased flight activity compared to controls that received non-infectious blood meal. Interestingly, this altered behavior persists for three weeks covering the entire extrinsic incubation period as well as the first week of the infectious stage (**Fig. 1c,d**). The increased flight activity of *An. stephensi*-*Pf* NF54 is mirrored by increased sugar feeding (**Fig. 1f**). None of the other mosquito-pathogen combinations showed major changes in flight statistics or sugar feeding behavior (**Fig. S2**). We analyzed population-level activity patterns and sugar feeding behavior at hourly intervals using generalized linear mixed models (GLMM). Consistent with the daily averaged results, the GLMM analysis shows that the increase in flight activity and sugar feeding observed in *An. stephensi*-*Pf* NF54 was most pronounced during the nocturnal active phase and across the light/dark transition (**Fig. 1g**). In contrast to *An. stephensi* infected with *Pf* NF54, we did not observe changes in the behavioral rhythm of *An. stephensi* infected with *P. falciparum* strain NF135 (*Pf* NF135, Asian origin) or *P. vivax* (Asian origin)*. An. gambiae s.s.* and *An. coluzzii*, both infected with *Pf* NF54, and *Ae. aegypti* infected with DENV also did not exhibit pathogen-associated changes in activity patterns, flight statistics, or sugar feeding behavior, suggesting that pathogen infections may have little impact on the basal activity patterns of mosquitoes across a variety of mosquito-pathogen combinations.

### Pathogen infection-induced changes in blood feeding behavior are species specific

Next, we asked if pathogen infection would impact blood feeding behavior in response to host-associated cues. To address this question, we adapted a previously developed experimental paradigm, the biteOscope, that uses a transparent artificial bite substrate including CO_2_ and heat as host cues, to visualize blood feeding mosquitoes in high resolution (**Fig. 2a**). As behavioral changes are most likely to have an impact on onward pathogen transmission during the infectious phase when the pathogen has disseminated to the salivary glands, we performed biteOscope experiments at 14 dpi. Taking advantage of the high-throughput nature of the biteOscope platform, we characterized the blood feeding behavior of 5113 mosquitoes in detail using matched infected and uninfected cohorts of 25 female mosquitoes per cohort for *Anopheles* experiments and a mean of 28 females for *Ae. aegypti* (see Table S2 and Materials and Methods for full details on experiments and replicates). All infected cohorts had an infection prevalence exceeding 90% (see **Tables S6** and **S7**). We developed a deep learning-based video analysis pipeline to quantitatively characterize short-range attraction, and exploratory, probing, and engorging behaviors in high throughput. Applying this pipeline to the full 5113 mosquito cohort yielded a robust dataset of over 98,000 landing events and ensuing behavioral trajectories of which 36,934 tracks were analyzed in depth using our deep learning based behavioral classification algorithm (trajectories shorter than 2 seconds were excluded from behavioral analysis, see **Fig. 2b** and Materials and Methods for full inclusion criteria).

**Figure 2.**
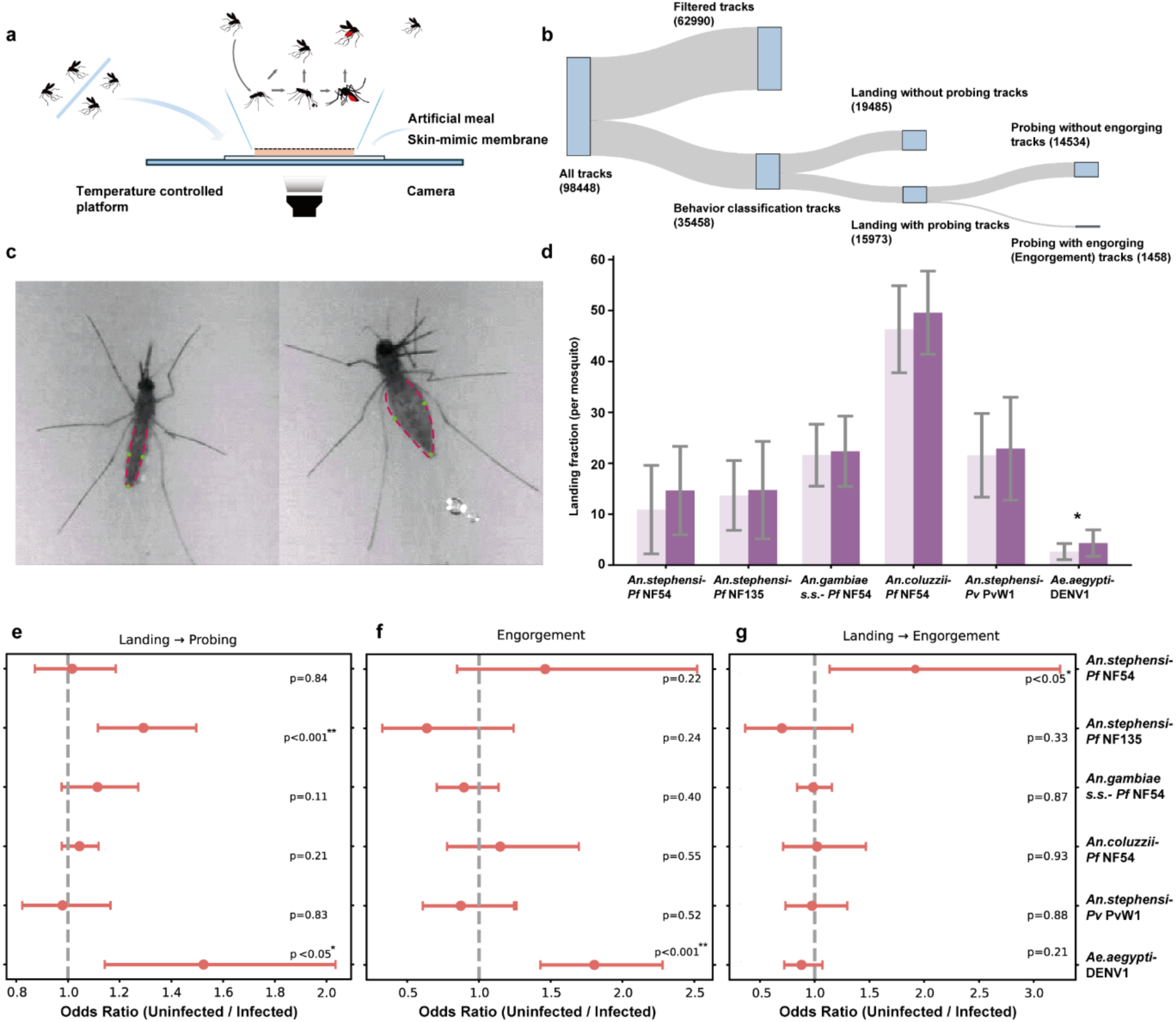
Cohort-level analysis of blood feeding behavior. **a.** Schematic of the biteOscope assay. An artificial blood meal covered by a membrane maintained at 37°C is mounted in the bottom of a cage. The bite substrate is transparent facilitating video recording of mosquitoes interacting with the substrate using a camera mounted below the cage. using pathogen-infected mosquitoes. The schematic depicts the different behavioral trajectories mosquitoes may engage in: upon landing, mosquitoes start exploring and subsequently may leave or pierce the surface and start probing. Upon probing, mosquitoes may leave, revert to exploratory behaviors, or engorge the artificial meal. Deep-learning based behavioral classification enables automatic quantification of distinct behaviors. **b.** Sankey plot of all behavioral trajectories. Each node represents the number of tracks at each processing step. **c.** An *An. stephensi* mosquito engorging on the biteOscope artificial blood meal. Clearly visible changes in abdominal morphology are used for automated computational detection of engorgement. **d.** Per mosquito landing rate of uninfected (light purple, left) and infected (dark purple, right) mosquitoes across different mosquito-pathogen combinations. Bars represent mean landing rate with error bars representing standard deviation. Statistical significance was assessed using the Wilcoxon rank-sum test (*p < 0.05). **e.** Forest plot of probing behavior following landing in infected versus uninfected mosquitoes across different mosquito-pathogen combinations. Points represent odds ratios for each mosquito-pathogen combination, calculated as the number of trajectories in which mosquitoes probed after landing, relative to landing events that did not lead to probing. Horizontal lines in panels e-g indicating 95% confidence intervals, the vertical dashed line denotes no effect (OR = 1), *p*-values from Fisher’s exact tests are shown for each comparison. **f.** Forest plot of the odds ratio of engorgement of infected versus uninfected mosquitoes across different mosquito-pathogen combinations for each mosquito-pathogen combination. **g.** Forest plot of the odds ratio of engorgement following landing in infected versus uninfected mosquitoes across different mosquito-pathogen combinations.

Following the behavioral sequence leading to a blood meal, we start our analysis by comparing landing rates of infected and uninfected mosquitoes. We defined each landing event as a successful host-seeking event and observed a variety of behaviors: Some mosquitoes engorge immediately upon first contact, others land multiple times prior to engorging, while other mosquitoes may not engorge at all despite multiple contacts with the host mimic. For all five *Anopheles-Plasmodium* combinations, we observed no differences in landing rate between infected and control groups, suggesting that *Plasmodium* infection has no or very limited impact on short-range host seeking behavior. In contrast, DENV infected *Ae. aegypti* exhibited a 1.6-fold increased landing rate (*p* < 0.05, Wilcoxon rank-sum test, see **Fig. 2d**) compared to uninfected controls, suggesting DENV enhances *Ae. aegypti’s* drive to seek blood. This observation is in line with a previous study reporting increased short-range host seeking behavior in DENV infected *Ae. aegypti* ^32^.

Upon landing, a mosquito may probe the skin surface in search of a blood meal. As this process is an efficient means of pathogen transmission, also when it does not lead to the uptake of blood, we asked if the propensity of mosquitoes to engage in probing behavior was subject to infection status. To address this question, we took advantage of the fact that the abdominal swelling resulting from engorgement is clearly visible in the biteOscope setup, and identified landing events leading to successful engorgement by analyzing temporal changes in abdominal morphology (**Fig. 2c**). Next, we separated landings in events that did from those that did not lead to engorgement. Focusing on landings that did not lead to engorgement, we observed that both *An. stephensi* infected with *Pf* NF135 and *Ae. aegypti* infected with DENV were more likely to engage in probing behavior and subsequently leave the bite substrate without engorging, compared to their uninfected counterparts (*An. stephensi*-*Pf* NF135: OR=1.29, *p* < 0.001; *Ae. aegypti*-DENV: OR=1.52, *p* < 0.05, Fisher exact test, see **Fig2e** and **Table S2**). This observation may have important implications for pathogen transmission, as probing alone (without engorgement) is generally sufficient for pathogen transmission to a vertebrate host ^32,42^. Furthermore, as probing yet non-engorging mosquitoes will not be satiated, they may engage in additional blood feeding attempts, thereby potentially stimulating onward transmission.

Following successful landing and probing, a mosquito may start blood engorgement until serum-derived compounds and abdominal distention signal completion^43^. We analyzed the fraction of mosquitoes that engorged per experimental cohort for infected and uninfected mosquitoes and observed a significantly increased engorgement rate for *Ae. aegypti* mosquitoes infected with DENV (OR = 1.77, *p* < 0.001, Fisher’s exact test). This enhanced tendency to take up a blood meal is likely caused by the increased landing rate of DENV infected *Ae. aegypti* (the odds to engorge after landing did not differ between infected and control cohorts, *p* = 0.21 Fisher’s exact test). Strikingly, all 5 *Anopheles-Plasmodium* combinations showed no effect of infection on the population fraction that took up a blood meal (**Fig. 2f**). In addition to population level engorgement statistics, we also analyzed the likelihood to engorge relative to all landing events. In this context, we observed a significant increase for *An. stephensi* infected with *Pf* NF54 (*p* < 0.05, Fisher’s exact test, **Fig. 2g**), indicating that upon landing, *Pf* NF54 infected *An. stephensi* are more likely to engorge compared to uninfected *An. stephensi*. Collectively, these results indicate that DENV induces marked alterations in *Ae. aegypti* blood feeding behavior by enhancing short range host seeking, probing, and engorgement behaviors. The *Plasmodium* infected anophelines show a more complex picture in which *An. stephensi*’s behavior seems to undergo alterations by some, yet not all parasites tested. Both *P. falciparum* lines enhanced aspects of biting behavior, with *Pf* NF153 enhancing probing and *Pf* NF54 increasing the likelihood to engorge upon landing, while *P. vivax* infection did not affect any of the behavioral parameters tested. Remarkably, the short-range attraction, biting, and engorgement dynamics of both *An. gambiae s.s.* and *An. coluzzii* remained unaffected by *P. falciparum* infection. The behavioral robustness of these two important African malaria vectors in the context of *P. falciparum* infection, underscores that pathogen-induced behavioral changes are not a general theme applicable to all mosquito-pathogen systems yet may differ markedly per mosquito and pathogen species.

### Fine-scale behavioral dynamics of blood feeding trajectories show diverse and species-specific patterns in pathogen-infected mosquitoes

Having analyzed short-range host seeking and blood feeding behaviors at the cohort level, we next asked if the detailed behavioral trajectories leading up to the blood meal differed between infected and uninfected mosquitoes. Complementing populational-level metrics, trajectory-level behavioral statistics capture fine-scale behavioral parameters (see **Fig. 3a**), and may reveal subtle effects of infection on behavior not apparent in coarse-grained population statistics. Analogous to our analysis above, we analyze behavioral trajectories that did, or did not, lead to engorgement separately. Notably, we observed that engorging *An. stephensi* infected with *Pf* NF54 spend approximately 150 seconds longer on the bite substrate, compared to control mosquitoes (*p* < 0.05, Wilcoxon rank-sum test) (**Fig. 3b,c**).

**Figure 3.**
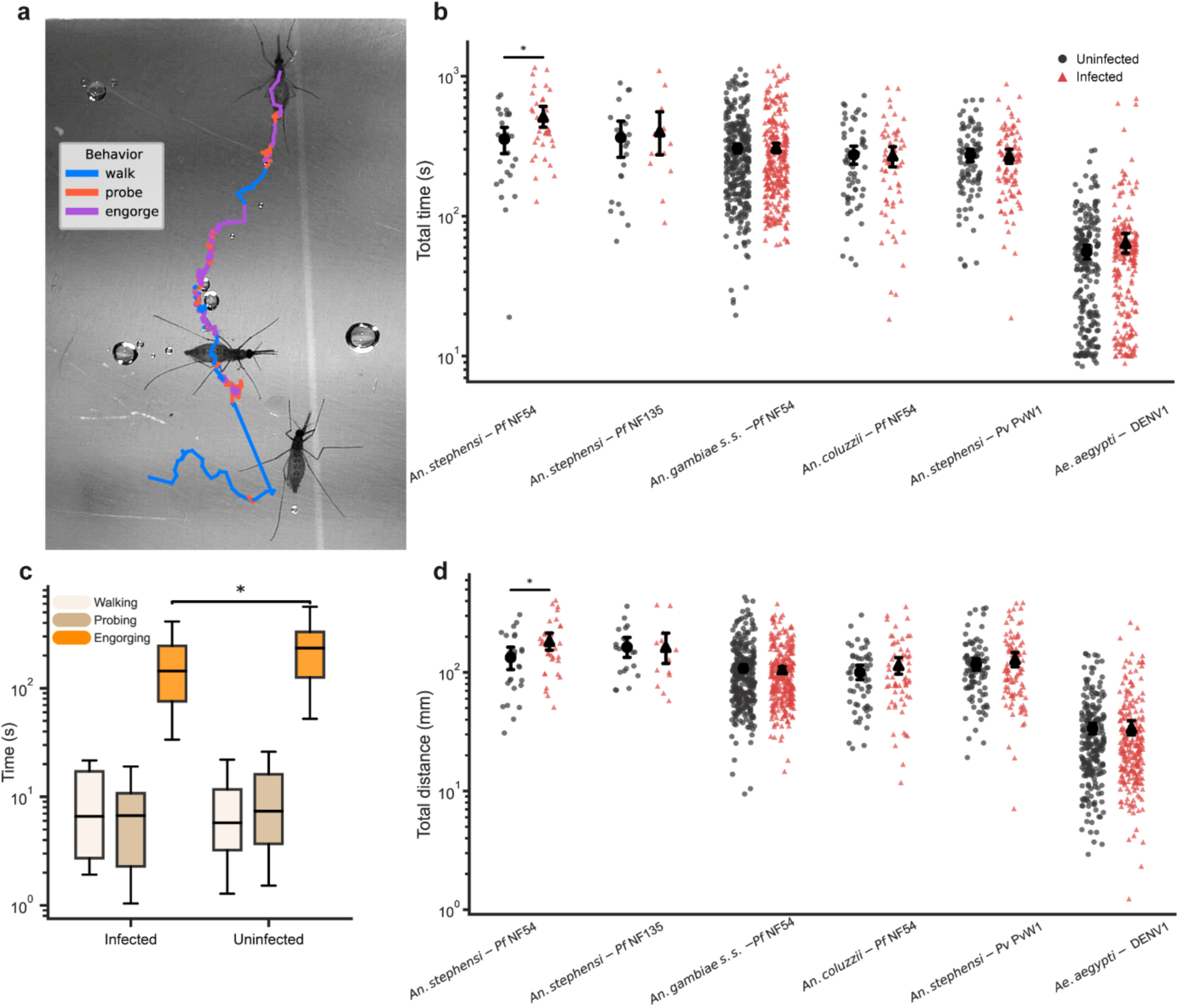
Trajectory-level analysis of biting and engorging behavior. **a.** A representative track showing movement of an individual mosquito (visible at the top of the frame) over the artificial bite substrate. Colors represent distinct behaviors (walking/exploration, probing, and engorging) as classified by deep-learning. The mosquito landed (bottom frame) and explored the bite substrate (blue section) to subsequently engage in several probing and engorging bouts. At the end of the track the mosquito is fully engorged as evidenced by the swollen abdomen **b.** Total time spent on the bite substrate across six mosquito–pathogen combinations. Each datapoint is derived from an individual trajectory of an engorging mosquito, error bars indicate standard deviation, statistical significance was assessed using the Wilcoxon rank-sum test (\**p* < 0.05). **c.** Behavioral decomposition of tracks leading to engorgement into walking, probing, and engorging time with different colors representing distinct behaviors. The observed difference in total time between *An. stephensi* infected with Pf NF54 and controls was primarily attributable to variation in engorging time. Error bars indicate standard deviation, statistical significance was assessed using the Wilcoxon rank-sum test (\**p* < 0.05). **d.** Total distance walked on the skin mimic across six mosquito–pathogen combinations. Each datapoint is derived from an individual trajectory of an engorged mosquito, error bars indicate standard deviation, statistical significance was assessed using the Wilcoxon rank-sum test (\**p* < 0.05).

Using our deep learning behavioral classification tool, we next decomposed the behavioral trajectories leading to engorgement into exploratory/walking, probing, and engorging behaviors. The increased residency time observed in *An. stephensi* infected with *Pf* NF54 was primarily attributable to a prolonged engorgement time, while the duration of exploring and probing was not influenced (**Fig. 3c**). The distance that mosquitoes walked on the biting surface—representing exploratory behavior—also increased in *An. stephensi* infected with *Pf* NF54. In a natural context, the prolonged engorgement time and increased walking distance may make *Pf* NF54 infected *An. stephensi* more vulnerable to host defensive behaviors potentially increasing mortality. These infection-associated changes in engorgement behavior seem to be particular to *Pf* NF54, as *Pf* NF135 and *P. vivax* infection did not result in similar behavior alterations (**Fig. 3b,d**). Mirroring population-level metrics, the fine-scale characteristics of engorging behavior of *An. gambiae s.s.* and *An. coluzzii*, did not differ between infected and control mosquitoes. Compared to all anophelines, engorging *Ae. aegypti* exhibited a substantially shorter residency time (∼60 seconds) on the biting substrate, while the walking distance covered during exploratory behavior was similar to the distance covered by *Anopheles* mosquitoes. DENV infection did not alter these behavioral parameters. Our behavioral classification analysis allows quantitative assessment of a variety of additional behavioral metrics, including the fraction of time spent moving, walking velocity, and morphological features of the mosquito’s abdomen serving as a proxy for the obtained meal size (maximum abdominal width, abdominal increase). We observed no differences between infected and control mosquitoes for any of these metrics for all six mosquito-pathogen combinations investigated (see **Fig. S6-S8** and **Table S3**).

Following our behavioral analysis of engorgement, we next turned our attention to individuals that landed on the bite substrate yet did not engorge. We observed that mosquitoes often landed multiple times without engorging, resulting in a vast dataset of 35,458 behavioral trajectories. Having observed an increased propensity for transitioning to probing after landing for *Pf* NF135 infected *An. stephensi* and DENV infected *Ae. aegypti* in our population level analysis, we next asked if infection would also have an effect on the number of probing bouts along a behavioral trajectory, or the duration of probing. For landing events not leading to engorgement, we indeed observed an increased number of probing bouts in both *An. stephensi-Pf* NF135 and *Ae. aegypti*-DENV (*p* < 0.05). Interestingly, the total duration of probing was longer in *Pf* NF135 infected *An. stephensi* compared to controls (7.3 vs. 6.4 seconds, *p* < 0.05). Longer probing times were also observed in *Pf* NF54 infected *An. stephensi*, while DENV infected *Ae. aegypti* showed the opposite trend: DENV-infected mosquitoes probed an average of 10.3 seconds compared to 14.0 seconds for uninfected controls (*p* = 0.05). The remaining three mosquito-pathogen combinations showed no differences in the number of probing bouts or probing duration (**Fig S9**). Infection furthermore showed a variety of significant, yet subtle effects on locomotion parameters, such as the time spent moving versus stationary (all *Anopheles-Plasmodium* combinations). While statistically significant, the modest effect sizes observed regarding locomotion statistics leads us to conclude that the impact of infection on the locomotion behaviors we report is unlikely to meaningfully impact the transmission biology of mosquito-borne pathogens, in contrast to the changes observed in probing and engorging behaviors. To provide further context regarding the relevance of our observations for pathogen transmission, we explore the effect of infection-induced behavioral changes on pathogen transmission using a conceptual epidemiological model below.

### Pathogen-induced behavioral alterations enhance the transmission of mosquito-borne pathogens

Taking a comparative perspective across mosquito-pathogen combinations, our results reveal that different aspects of blood feeding behavior are subject to change upon infection, depending on the mosquito-pathogen combination in question. We explore the impact of the observed behavioral alterations on the transmission of the respective pathogen, using a conceptual mathematical model for mosquito-borne pathogens (see Supplementary Information and Fig. S10). We conceptualize the epidemiological dynamics^44,45^ using a compartmental model in which infection dynamics are governed by interactions between human hosts and mosquito vectors. An important parameter in this model is the interaction rate of mosquitoes and humans (typically referred to as the "biting rate"). While blood feeding is a complex behavioral routine, models often have a single biting rate parameter, *B_i_,* as a composite metric abstracting the entire blood feeding process into a single parameter. Using this model we gained a conceptual understanding of the impact of behavioral changes on pathogen transmission by estimating the maximum value of the time-varying effective reproductive number (𝑅𝑡_𝑚𝑎𝑥_) of the pathogen while varying *B_i_* (**Fig. 4**). Our experiments revealed an increased likelihood to probe after landing for both *Pf* NF135 infected *An. stephensi* (OR = 1.3) and DENV infected *Ae. aegypti* (OR = 1.5). Interpreting this enhanced tendency to probe as an increased biting rate in our model, translates to a 1.4 fold increase in 𝑅𝑡_𝑚𝑎𝑥_. We also observed higher engorgement rates (*Ae. aegypti*-DENV OR = 1.8) and an increased likelihood to engorge after landing (*An. stephensi*-Pf NF54, OR = 1.9). Interestingly, a doubling of the engorgement, and therefore biting rate, nearly results in a tripling of *R* illustrating that relatively minor variations in biting rate correspond to large increases in *R* due to non-linear effects.

**Figure 4.**
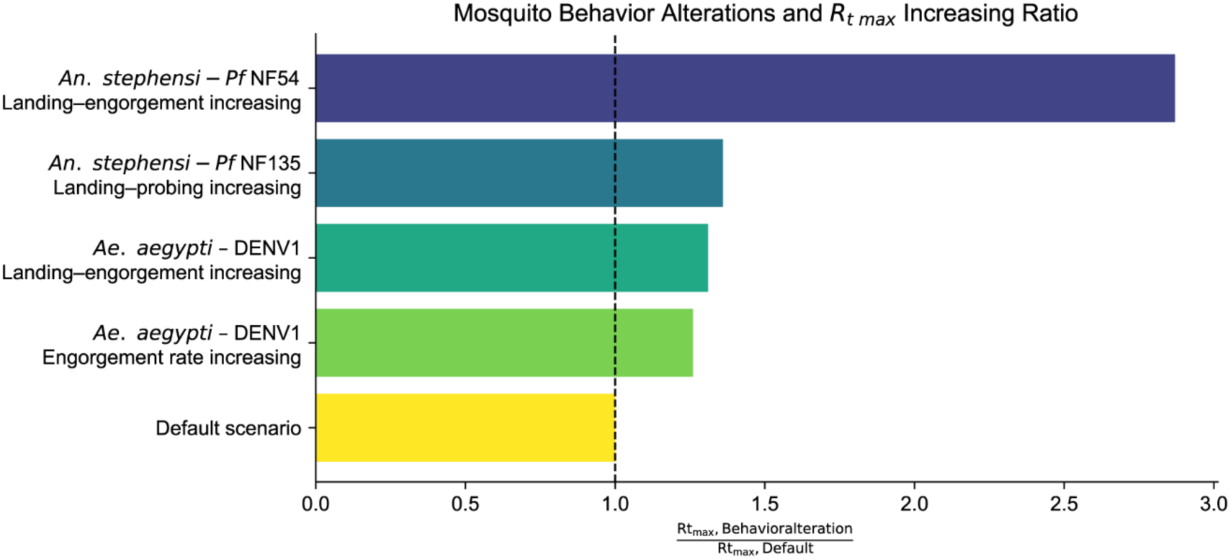
Modelling the impact of infection-induced behavioral changes in mosquito host-seeking and biting behavior on pathogen transmission. Ratio of the maximum value of the time-varying effective reproductive number between mosquitos exhibiting infection-associated change in biting rate. Each bar represents the peak transmission potential corresponding to a specific behavioral alteration pattern.

## Discussion

We combined high-resolution video tracking and deep-learning-based computational analysis to quantify the impact of pathogen infection on mosquito behavior across six mosquito-pathogen combinations of high public health importance. Utilizing two powerful behavioral assays allowed us to characterize diverse aspects of mosquito behavior during progression of the infection, and obtain rich behavioral data at the peak of infectiousness. Our comparative approach using identical assays for different mosquito-pathogen combinations provides a unique perspective revealing clear species-specific effects.

We extensively studied the originally Asian malaria vector *An. stephensi* that has recently invaded the Horn of Africa^46,47^. Interestingly, the behavior of this mosquito species was affected most by *P. falciparum* NF54, a parasite line originally isolated from Africa. *Pf* NF54 increased *An. stephensi*’s flight activity starting from the early phases of infection, and this effect lasted throughout the entire three week monitoring period. These effects were not observed upon infection by *P. falciparum* NF135, a *P. falciparum* line originating from Asia, nor by *P. vivax* which is also common in the native range of *An. stephensi*. Dissecting biting behavior during the period of peak transmission, we also observed behavioral differences in relation to *Pf* NF54 infection in *An. stephensi* resulting in an enhanced tendency to probe after landing and a significant increase in engorgement time. Again, these effects were not observed in the context of *P. vivax* or *Pf* NF135 infection. Together, these data give credence to the previously voiced hypothesis^21,22^ that parasites infecting sympatric mosquito species have limited impact on mosquito behavior due to a shared evolutionary history having attenuating effects on the impact of infection. In contrast, parasites infecting mosquitoes originating from non-overlapping geographical ranges that do not share an evolutionary history may have a larger impact on behavior. This hypothesis is further supported by our observation that the African *P. falciparum* strain (NF54) had no significant effect on the behavior of the African vectors *An. gambiae s.s.* and *An. coluzzii*.

Our results concerning DENV infected *Ae. aegypti* paint a different picture. Both the mosquito colony and DENV strain used are relatively recent isolates from the same region (Thailand). While we observed no impact of infection on flight activity patterns along the three week infection period, we observed significant differences in blood feeding behavior between infected and uninfected *Ae. aegypti*. At 14 dpi, DENV infected *Ae. aegypti* more readily engaged in short-range host-seeking behavior, exhibited a higher probing frequency, and a large fraction of DENV-infected mosquitoes took a blood meal– showing a clear impact of infection on behavior. These observations are in line with a previous study by Xiang *et al.* reporting increased short-range host seeking and probing without subsequent engorgement, although Xiang *et al.* did not observe a difference in the number of mosquitoes that engorged, likely due to a much smaller number of mosquitoes included in the study^32^. The increased tendency to probe but not take a blood meal that DENV infected mosquitoes exhibit is likely to induce additional blood feeding attempts, as observed previously by Maciel-de-Freitas *et al.*^28^ , leading to additional bites and thus likely enhancing DENV transmission. Interestingly, Xiang *et al.* and Maciel-de-Freitas *et al.* used a DENV strain of serotype 2, whereas we used serotype 1 suggesting similar effects on biting behavior of these serotypes.

Building upon our previously developed high-resolution and -throughput behavioral assays, we were able to quantify the effects of pathogen infection on mosquito behavior at an unprecedented scale, resulting in statistically robust estimates of infection-associated behavioral change. Importantly, applying this approach to six mosquito-pathogen combinations allows us to conclude that behavioral alterations are not universal features of mosquito–pathogen interactions as is often assumed, but can be highly species- and strain-specific. The behavioral effects we observed were generally subtle, and may have remained undetected in smaller-scale studies; however, the scale and precision afforded by these assays allowed us to uncover consistent shifts in activity and blood-feeding behavior. Although the behavioral effects we identified were modest in magnitude, our conceptual transmission model demonstrates that even small changes in host-seeking and feeding behavior can substantially amplify onward pathogen transmission, underscoring the potential epidemiological importance of these subtle phenotypic shifts.

## Materials and Methods

All mosquito and pathogen species used in this study are detailed in **Table S1**.

### Mosquito rearing

Mosquitoes were reared according to the following procedures.

*An. stephensi, An. coluzzii, An. gambiae s.s.*: Eggs were introduced in larval trays filled with tap water and maintained at ∼200 larvae/liter in a climate-controlled climate (27–30°C and 70-80% humidity), fed on fish food Liquifry (Liquifry No1, Interpet) and TetraMin baby (Tetra). Adult mosquitoes were collected by aspiration and transferred to netted cages. Female and male mosquitoes were not separated and maintained in an insectary (28–30°C and 70-80% humidity) with a 12-hour day/night cycle including 30 minute dimming transitions. Mosquitoes had continuous access to 5% glucose mixed with 0.4% methylparaben.

*Ae. aegypti*: Eggs were hatched in a vacuum chamber for 45 minutes. Approximately 200 larvae were transferred per larvae tray filled with 1.5 liter tap water, feeding on TetraMin baby (Tetramin, Tetra). Adult *Ae. aegypti* were maintained in an insectary of 28 ± 1°C, 75% relative humidity with a 12-hour day/night cycle including dimming transitions, having continuous access to 10% sucrose.

### *P. falciparum in vitro* culture and mosquito infection

*P. falciparum* strain NF54 (West Africa)^48,49^ and NF135 (Cambodia)^50^ were cultured *in vitro* as previously described^51^. Briefly, *P. falciparum* gametocytes were continuously cultured in RPMI medium (RPMI1640; Life Technology Invitrogen, 518-00035) mixed with erythrocytes and serum from malaria naïve donors (Sanquin; Nijmegen, The Netherlands). A semi-automated tipper system was adapted for culturing at 37 °C, and continuously supplied with a pre-mixed gas mixture of 93% N_2_, 3% O_2_ and 4% CO_2_. At day 14 after seeding, gametocytes were collected and activated to assess the exflagellation ability of male gametocytes by the presence of exflagellation centers in fetal bovine serum (Invitrogen, 10270106). Cultures that exflagellated were used to prepare an infectious blood meal.

Approximately 300 female *Anopheles* mosquitoes (1 to 3-days-old) were collected and secured in two netted cages (150 females each). Hundred fifty female mosquitoes were provided with an infectious blood meal prepared with gametocyte culturing material mixed with human blood, using midi-feeders (1.2 ml) maintained at 37 °C. A second batch of 150 female mosquitoes was provided with a control blood meal not containing parasites. Feedings took place for 30 minutes in darkness. After feeding, unfed and partially fed mosquitoes were removed. Forty fully-fed mosquitoes were directly transferred to the BuzzWatch assay. All remaining mosquitoes, to be used for biteOscope assays at 14 dpi, were maintained in netted cages with continuous access to 5% glucose. On day 7 after receiving the initial blood meal, both infected and control mosquitoes were offered a second, non-infectious blood meal.

### *P. vivax* mosquito infection

A cryopreserved stabilate of red blood cell (RBC) infected with *P. vivax* strain PvW1 (Thailand)^51^ was obtained from University of Oxford (Oxford, UK). Infected RBC samples were thawed and used to inoculate malaria-naïve, healthy adult study participants who were Duffy blood group-positive (Registered under the study ID NL-OMON57011). Methods for the preservation of infected RBC and blood inoculation were previously described^51^. Between day 11 to day 25, peripheral blood was repeatedly sampled to assess transmission of *P. vivax* to *An. stephensi*. Samples with gametocyte numbers that consistently led to at least 80% infected mosquitoes were used in this study. Forty fully engorged mosquitoes were directly used for the BuzzWatch experiments, while all remaining mosquitoes were maintained in netted cages, receiving a second non-infectious blood feeding 6 days post first blood meal.

### Determination of mosquito *Plasmodium* infection rate

To determine infection prevalence and load, a subset of *Anopheles* mosquitoes that received a *P. falciparum-* or *P. vivax*-infected blood meal were dissected to determine the infection prevalence and load at 7 dpi. Briefly, a minimum of 20 mosquitoes were taken from each cage, dissected for midguts and stained with 1% mercurochrome to assess the presence of oocysts. Cages with oocyst infection prevalence exceeding 80% were kept for experiments. On one occasion, a BuzzWatch experiment was terminated because an *An. gambiae s.s.* cohort did not meet the >80% oocyst infection prevalence criteria, the corresponding results were excluded from final analysis.

After completion of behavior assays, cages were frozen at -20°C to kill mosquitoes. Subsequently, *Plasmodium* infection prevalence in each experimental batch was determined by qPCR following a previously described protocol^53^. For each assay, 20 mosquitoes were randomly selected from each BuzzWatch cage and 10 mosquitoes from each biteOscope cage, and individually placed in separate wells of 96-well plates (CoStar, 3958), containing 0.2 g of 1-mm zirconia beads (Biospec, 11079110z) and 100 µL of PBS, beaten for 10s (Biospec, Mini Beadbeater 96) and centrifuged at 2000 rpm for 1 min. Supernatant was collected and mixed with 150 µL PBS. Total DNA was extracted and eluted in 100 μL with the automated MagNaPure LC instrument (MagNaPure LC DNA Isolation kit, Roche). The presence of *P. falciparum* and *P. vivax* was determined by qPCR targeting the 18S rRNA small subunit gene, with primer and probe sequences adapted from previous reports ^54,55^ on a CFX96 Real-Time PCR Detection System (Bio-Rad). Each qPCR run included positive (pure sporozoite extraction) and no-template negative controls (mosquito receiving non-infectious blood meal, and PBS only) to validate amplification specificity and exclude contamination.

### DENV-1 virus preparation and mosquito infection

Viral stocks of DENV-1 isolate KDH0026A (Thailand) **(Table S1)** were prepared using *Aedes albopictus* C6/36 cells as previously described^56^. In brief, C6/36 cell monolayers at a sub-confluent stage were inoculated with 0.2 mL of viral suspension and incubated for 7 days at 28 °C. Cells were maintained in Leibovitz’s L-15 medium containing penicillin, streptomycin, non-essential amino acids, and 2% fetal bovine serum (Life Technologies). After the incubation period, culture supernatants were harvested, supplemented with 10% FBS, adjusted to a pH of ∼8 with sodium bicarbonate and stored at -80 °C.

Prior to receiving a blood meal, 5-7 day old female mosquitoes were collected and starved for 24 hours. Mosquitoes were offered an infectious blood meal for 15 minutes using a membrane-feeding system (Hemotek). The artificial meal was prepared as a 2:1 mixture of washed human erythrocytes (Institut Pasteur, Paris) and virus suspension at a final concentration between 1.25 x 10^6^ and 4.5 x 10^6^ focus-forming units (FFU)/ml, supplemented with 10mM adenosine triphosphate. Fully engorged mosquitoes were then either directly transferred into the BuzzWatch cages in groups of 40 per cage, or sorted into BiteOscope cages and kept in incubators set at 28 ± 1°C, 75% relative humidity with a 12-hour day/night cycle (including dimming transitions), with continuous access to 10% sucrose.

### Determination of DENV infection rate in mosquito

After BiteOscope experiments, mosquitoes were knocked-down on ice and taken out from cages and homogenized individually in 400 μl of RAV1 RNA extraction buffer (Macherey-Nagel, 740928.40) during two rounds of 30 sec at 5,000 rpm in a TissueLyser II grinder (Qiagen). Total RNA was extracted using the NucleoSpin 96 kit (Macherey-Nagel, 740709.4) following the manufacturer’s instructions. Total RNA was converted to complementary DNA (cDNA) using M-MLV reverse transcriptase (Invitrogen, 28025013) and random hexamers. The cDNAs were amplified by PCR. The amount of DENV-1 NS5 was measured by qPCR using the QuantiTect SYBR Green kit (Qiagen, 204143) on a LightCycler 96 real-time thermocycler (Roche) following a published protocol ^57,58^. The *Ae. aegypti* ribosomal protein-coding gene RP49 (AAEL003396) was used for normalization ^59^. The relative cDNA quantity was calculated as E-(CqRP49-Cq16S), with E being the PCR efficiency of each primer pair. For Buzzwatch experiments: after feeding mosquitoes a DENV infectious blood-meal, 40 fully engorged females were placed in the BuzzWatch cages. A second group of 40 fully engorged females were placed in paper boxes supplemented with sugar cotton and kept in a climatic chamber identical to the BuzzWatch set-ups. The second group was used to determine infection prevalence of DENV-1 at 14 dpi by PCR. cDNA of whole mosquitoes was obtained similarly to the BiteOscope experiments (see above) and the cDNA was subsequently amplified using DreamTaq DNA polymerase (Thermo Fisher Scientific) and specific primer pairs for DENV-1 (using primers P17 P18 and methods from^60^). Amplicons were visualized by electrophoresis on 2% agarose gels.

### Long-term activity monitoring - BuzzWatch assay

For longitudinal monitoring of mosquito behavior, we adapted the BuzzWatch assay^35^. The design and construction of the setup is available at: https://theomaire.github.io/buzzwatch/construct.html. A Raspberry Pi V2 NOIR 8 MP camera covered with an 850 nm infrared long-pass filter (Thorlabs, FEL0850) was placed at the bottom of the cage to image mosquitoes using infrared illumination. Each camera was connected to a Raspberry Pi 4B+ microcomputer configured to automatically capture video in h264 format. Recordings were saved as 20-minute segmented videos (25 frames per second) and converted to mp4 format every 12 hours. Prior to the experiment, recording parameters were manually adjusted via the Python *pirecorder* module^61^ to maximize recording quality.

After blood feeding, 40 fully engorged mosquitoes were aspirated into their corresponding cages. Two sugar feeders filled with cotton were attached to opposite sides of the cage, with *Anopheles* receiving 5% glucose and *Aedes* 10% sucrose. Cages were moved into an enclosure placed in a climate chamber (28–30°C and 70-80% humidity). Two RGB LED strips were mounted inside the enclosure and controlled by a Raspberry Pi to reproduce diurnal light cycles consisting of 12 h of darkness and 12 h of light with gradual transitions at artificial dawn and dusk. Environmental parameters such as temperature and humidity were continuously monitored by a Raspberry Pi. Each BuzzWatch experiment lasted for 3 weeks (0--21 dpi). Every 7 days, the recording was paused for 30 minutes to replace the cotton in the sugar feeders. On day 22 post infection, all mosquitoes were cold anesthetized and moved out of the cages, and stored at -80°C for qPCR-based infection rate determination.

### BuzzWatch data analysis

The mosquito tracking pipeline followed our previously developed approach^35^. Briefly, standard computer vision methods were applied to separate mosquitoes from the cage background. The reference background was generated by averaging 100 frames across 100 consecutive 20-minute videos. Resting mosquitoes were then detected using a blob detection algorithm, followed by the greedy algorithm generating continuous tracks. Flying mosquitoes were detected by applying optimized segmentation and centroid tracking after inter-frame subtractions. A custom algorithm was developed to link the tracks of resting and flying mosquitoes, allowing the identification of behavioral state transition between take-off and landing. This algorithm performs a local search on spatially adjacent track endpoints to find matching pairs. The assumption of take-off events was based on the spatial proximity between the endpoint of a resting track and the starting point of a flying track, and conversely for landings. After processing, the final tracking result including mosquito IDs and corresponding position coordinates were saved for behavioral quantification.

We analyzed data from each mosquito-pathogen combination separately, with 2-4 experimental replicates per combination. All time-series data were resampled to 1-minute intervals and normalized by the number of alive mosquitoes and smoothed using a 20-minute rolling average. All downstream behavioral metrics and statistical analyses were computed based on these normalized and smoothed time series. The fraction of flying mosquitoes was calculated as the ratio of detected flying individuals and the total number of alive mosquitoes present in the cage at each time point. The sugar feeding index was calculated by comparing the fraction of mosquitoes resting on the sugar feeder surface to that on the control surface, with the control surface manually designated from the side of the cage where sugar feeders were not placed (**Figure S1**). Behavioral variables were averaged within hourly intervals for each experimental day. We used an interval-based GLMM analysis as described^35^, to report Z-scores derived from the GLMM analysis, indicating the strength and significance of pathogen infection effect on all behavioral statistics (fraction of flying mosquitoes, sugar feeding index, flight rhythm, flight speed, and flight duration).

### Fine-scale mosquito behavior recording - BiteOscope assay

Mosquito short range host-seeking and biting behaviors were recorded with BiteOscope assay^33,34^. BiteOscope cage design and construction can be found at https://github.com/felixhol/biteOscope/tree/master/cageDesigns. For experiments with *Anopheles* mosquitoes, mosquitoes were aspirated from netted cages into the biteOscope cages at 14 dpi, with an exact number of 25 mosquitoes per cage. For *Aedes* experiments, mosquitoes were directly introduced into biteOscope cages after receiving the infectious or control blood meal, with a maximum number of 35 mosquitoes per cage. All biteOscope experiments were conducted at 14 dpi. Prior to each experiment, mosquitoes were starved by providing only water for at least 4 hours. Each cage was then transferred onto an individual artificial bite substrate, which was created using a flat acrylic sheet heated to ∼37°C. A layer of Parafilm (M laboratory film) was tightly stretched across the plate to serve as the skin mimic. An artificial meal, consisting of 10 mL of solution containing 1 mM ATP (Sigma-Aldrich, A3377), 110 mM NaCl (Sigma-Aldrich, S7653), and 20 mM NaHCO₃ (Sigma-Aldrich, 31437-M), was introduced into the space between the Parafilm membrane and the heated acrylic sheet. When the artificial meal reached ∼37°C, the biteOscope cage was placed on top of the artificial meal. BiteOscope cages have a window in the bottom that is covered with a trapdoor that can be opened to give the mosquitoes access to the artificial bite substrate. The experiment was started by opening the trapdoor and providing a ∼10s pulse of human breath to the mosquitoes to stimulate blood-feeding behavior. A camera (Basler ace acA2040-90um), fitted with a 35 mm lens (Thorlabs MVL35M23) and controlled by pylon 5 software, was situated under the acrylic sheet to record 20 minute videos at 25 frames per second. *Anopheles* experiments were conducted in darkness using infrared illumination while *Aedes* experiments were conducted under daylight conditions. At the end of the recording, all mosquitoes were cold shocked and moved out of the cages, stored at -80°C for qPCR-based infection rate determination.

### BiteOscope tracking, behavioral classification, and analysis

Tracking of mosquito movements on the skin mimic was conducted using a custom convolutional neural network trained using DeepLabCut^36,62^. Due to differences in anatomy and illumination conditions, separate models were trained for *Anopheles* and *Aedes* mosquitoes. For the *Anopheles* model, a total of 50 manually labeled frames containing more than 500 mosquitoes were annotated with 37 keypoints and used to train a DLCRNet_ms5 model. For the *Aedes* model, 350 manually labeled frames including more than 500 mosquitoes (35 keypoints) were used to train a Resnet_50 model. For the *Anopheles* model, the test error was 8.31 pixels (0.33mm) and train error 2.75 pixels (0.11mm) for the *Aedes* model, the test error was 3.06 pixels (0.12mm) and train error 2.78 pixels (0.11mm). Trained networks were used for mosquito tracking and pose-estimation on biteOscope videos.

DeepLabCut derived coordinates of all detected body parts for every mosquito in each frame was subsequently assembled into tracklets with a minimum length of 25 consecutive frames. We considered each tracklet as a trajectory of an independent behavioral episode performed by a mosquito, beginning with landing and surface exploration by walking, followed by possible interaction with artificial meal (probing and engorging), ending with mosquitoes’ departure (**Fig 2a**). For the classification of walking, probing, and engorging behaviors, we applied a custom deep learning framework specifically developed for mosquito behavioral classification. This pipeline integrates manual video annotation, pose-based keypoint processing, and sequence classification into a unified workflow. Following manual annotation, pose data were normalized by translation, scaling, and rotation relative to reference keypoints to ensure spatial consistency across individuals. Sequential features were segmented into overlapping temporal windows of 48 frames and used as input to a convnet-based neural network^63^ designed to capture multi-scale temporal dynamics. The model was trained with grid search–optimized hyperparameters and evaluated using stratified cross-validation, with performance assessed by accuracy and F1-score. Considering morphological differences between *Aedes* and *Anopheles* mosquitoes as well as variations in video recording conditions such as lighting, independent models were trained for each species. During inference, only tracklets longer than 50 frames were included in the analysis. A behavioral state of walking or probing was assigned to a tracklet if at least one frame within it was predicted with a confidence score above 0.6, whereas frames with confidence below this threshold were excluded from subsequent quantitative analyses. For the classification of engorging behavior, we quantified temporal changes in abdominal width based on the distances between the left and right abdomen keypoints extracted from DeepLabCut tracking data. The raw abdominal width time series was smoothed using a rolling average of 25 frames to remove jitter. For each trajectory, maximum and minimum abdominal width were estimated as the 90th percentile of abdominal width along the whole trajectory of engorgement and the 10th percentile within a given time window at the start of the trajectory (400 frames, equivalent to 16 seconds for *Anopheles*; 200 frames, equivalent to 8 seconds for *Aedes*). The relative expansion of abdomen width was calculated as the ratio of the 90th percentile and the 10th percentile. A mosquito was classified as engorged when its abdominal width exhibited at least a 1.2-fold expansion. In addition, the 90th percentile of abdominal width was also required to exceed 18 pixels (0.72mm) for *Anopheles* and 30 pixels (1.20mm) for *Aedes*. All trajectories classified as engorged were subsequently verified by visual inspection of the corresponding video.

The individual-level behavioral statistics were derived from the characteristic of trajectory and the body parts coordinates. For each trajectory, the total residency time (Total time, s) was defined as the duration between the first and last frames. The time of specific behavior, namely walking (Walking time, s), probing (Probing time, s) and engorging (Engorging time, s), was defined as the total number of frames classified as that behavior by the behavioral classification framework. The total distance traveled on the biting substrate (Total distance, mm) was calculated as the smoothed cumulative displacement of thorax, head, and abdominal landmarks. The fraction of time spent moving (moving fraction) was defined as the ratio between the number of frames where thorax movement surpassed a set threshold (0.1mm) and the total number of valid frames in the trajectory. The mean speed (Mean speed, mm/s) was defined as the total distance traveled on the biting substrate divided by the total residency time.

### Malaria and dengue epidemiological modelling

We developed an SIR-SI (Susceptible-Infectious-Recovered human/Susceptible-Infectious mosquito) compartmental model to explore the potential impact of behavioral changes on *Plasmodium* and DENV transmission dynamics. The model is detailed below as a system of ordinary differential equations:

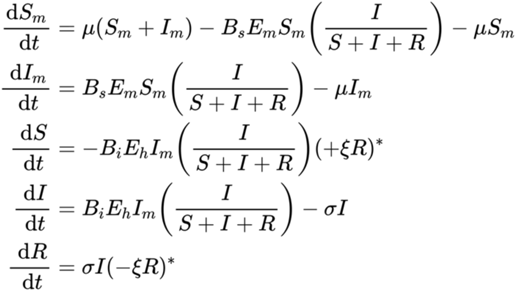

*\* ξR Only applicable for modelling malaria transmission*.

The meaning of each parameter and their baseline values used are described in detail in **Table S5**. The model includes susceptible and infectious mosquitoes, as well as susceptible, infectious, and recovered humans. We assume that at the beginning of simulation, the entire human population is susceptible. Infection happens only when a human interacts with a mosquito. Mosquito cannot naturally recover from infection. We also assumed that both human and mosquito population sizes were not influenced by infection. The following variables for both *Plasmodium* and DENV infection were set independently based on previous published studies (**See Table S5)** : Mosquito mortality rate (µ), susceptible (no-infection) mosquito biting rate (B_s_), the probability of mosquito infection upon exposure to an infectious human (E_m_), the probability of human infection following exposure to an infectious mosquito (E_h_), and the human recovery rate (δ). Specifically, to account for the time-dependent waning of host immunity in malaria patients^64,65^, an immunity loss constant (ξ) was incorporated into the *Plasmodium* model with the value derived from previous published studies. We ran the simulation for 2 years (730 days) with initial population sizes of 10000 mosquitoes, 500 susceptible humans, with 1 initial infectious human for both malaria and dengue fever scenario. To capture the long-term pathogen transmission dynamic driven by pathogen-induced mosquito behavioral alteration, we estimated the time-varying reproduction number (Rt), rather than the basic reproduction number (R0), as it can reflect changes in disease transmission potential over time. We applied R package deSolve^66^ for the epidemiological modelling, and R packages R0^67^ and EpiEstim^68^ for the calculation of Rt values.

The initial baseline model was constructed assuming mosquito behavior is not influenced by pathogen infection, by setting the biting rate of infectious mosquito to the same value as the biting rate of susceptible mosquito, 0.25. The maximum time-varying reproduction number (𝑅𝑡_𝑚𝑎𝑥_, 𝐷𝑒𝑓𝑎𝑢𝑙𝑡) of the model was captured to mark the transmission peak. We then incorporated behavioral alterations relating to mosquito probing and engorging behavior into the model by adjusting the infectious mosquito biting rate, as both *Plasmodium* and DENV can also be transmitted to their host through probing alone^69,70^. We captured the maximum time-varying reproduction number influenced by mosquito behavior alteration ( 𝑅𝑡*_max_*, 𝐵𝑒ℎ𝑎𝑣𝑖𝑜𝑟 𝑎𝑙𝑡𝑒𝑟𝑎𝑡𝑖𝑜𝑛 ) and estimated the 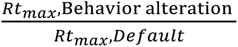 under five behavioral alteration scenarios:

1. Baseline scenario – uninfected mosquito behavior, with B_i_ set equal to 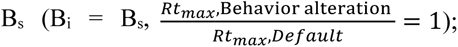
2. *An. stephensi*-Pf NF54 landing-engorging increasing – B_i_ multiplied by 1.92;
3. *An. stephensi*-Pf NF135 landing (no engorgement) – probing (no engorgement) increasing – B_i_ multiplied by 1.29;
4. *Ae. aegypti*-DENV landing (no engorgement) – probing (no engorgement) increasing – B_i_ multiplied by 1.52;
5. *Ae. aegypti*-DENV engorgement ratio increasing – B_i_ multiplied by 1.77.

All adjusted values were derived from the odds ratios obtained in populational-level behavioral statistical parameters from BiteOscope assay experiments (**Table S2**).

## Acknowledgments

We are grateful to Wouter Graumans, Nicolas Proellochs, Jolanda Klaassen, Laura Pelser-Posthumus, Astrid Pouwelsen, and Saskia Mulder (Radboud UMC) for breeding *Anopheles* mosquitoes; and Gregory Murray, Catherine Lallemand, Stéphanie Dabo, Sarah Merkling, Anna Beth Crist, and Fabien Aubry (Institut Pasteur) for help with *Aedes*-DENV experiments. We thank all the OptiViVax team members from University of Oxford, UK for their help with *P. vivax* experiments. We would also like to thank all team members from Louis Lambrechts lab at Institut Pasteur and Felix Hol lab at Radboud UMC for their discussion and feedback throughout this project, as well as their comments on the manuscript.

## Author contributions

Conceptualization: Z.W., T.M., E.G., T.B., B.M., L.L., and F.J.H.H. Software: V.K., Z.W., T.M. and F.J.H.H. Mosquito rearing and pathogen infection: E.G., T.M., R.S., and G.J.V.G. Experiments: Z.W., T.M., E.G., K.L., and F.J.H.H. Writing: Z.W., T.M. and F.J.H.H, with input from all authors.

## Ethics

The human sample involved in the *Plasmodium vivax* experiment in this study was under the OptiViVax project of Horizon Europe programme. The study was approved by the Central Committee on Research Involving Human Subjects (CCMO). All participants met the eligibility criteria (Supplemental Methods) and gave written informed consent before inclusion in the study. The trial was registered with the Overview of Medical research in the Netherlands (NL-OMON57011). Human blood samples to prepare mosquito artificial infectious blood meals at the Institut Pasteur were supplied by healthy adult volunteers at the ICAReB biobanking platform (BB-0033-00062/ICAReB platform/Institut Pasteur, Paris/BBMRI AO203/[BIORESOURCE]) of the Institut Pasteur in the CoSImmGen and Diagmicoll protocols, which had been approved by the French Ethical Committee Ile-de-France I. The Diagmicoll protocol was declared to the French Research Ministry under reference 343 DC 2008-68 COL 1. All adult subjects provided written informed consent.

## References

1. WHO. Vector-borne diseases. Vector-borne diseases https://www.who.int/news-room/fact-sheets/detail/vector-borne-diseases.

2. Coutinho-Abreu, I. V., Riffell, J. A. & Akbari, O. S. Human attractive cues and mosquito host-seeking behavior. Trends in Parasitology 38, 246–264 (2022).

3. van Breugel, F., Riffell, J., Fairhall, A. & Dickinson, M. H. Mosquitoes Use Vision to Associate Odor Plumes with Thermal Targets. Current Biology 25, 2123–2129 (2015).

4. McMeniman, C. J., Corfas, R. A., Matthews, B. J., Ritchie, S. A. & Vosshall, L. B. Multimodal Integration of Carbon Dioxide and Other Sensory Cues Drives Mosquito Attraction to Humans. Cell 156, 1060–1071 (2014).

5. McBride, C. S. et al. Evolution of mosquito preference for humans linked to an odorant receptor. Nature 515, 222–227 (2014).

6. Bansal, P., Pillai, R., Pooja, D. & Sen, S. Q. Two neuropeptides that promote blood feeding in Anopheles stephensi mosquitoes. Preprint at 10.1101/2024.05.15.594342 (2024).

7. Dittmer, J., Alafndi, A. & Gabrieli, P. Fat body–specific vitellogenin expression regulates host-seeking behaviour in the mosquito Aedes albopictus. PLoS Biol 17, e3000238 (2019).

8. Bento, I. et al. Parasite and vector circadian clocks mediate efficient malaria transmission. Nat Microbiol 10, 882–896 (2025).

9. Cator, L. J. et al. Immune response and insulin signalling alter mosquito feeding behaviour to enhance malaria transmission potential. Sci Rep 5, 11947 (2015).

10. Gaburro, J. et al. Neurotropism and behavioral changes associated with Zika infection in the vector Aedes aegypti. Emerging microbes & infections 7, 1–11 (2018).

11. Carrillo-Bustamante, P., Costa, G., Lampe, L. & Levashina, E. A. *Mosquito Metabolism Shapes Life-History Strategies of* Plasmodium *Parasites*. http://biorxiv.org/lookup/doi/10.1101/2022.07.06.498937<x> (2022) doi:10.1101/2022.07.06.498937.

12. Gomes, F. M., Silva, M., Molina-Cruz, A. & Barillas-Mury, C. Molecular mechanisms of insect immune memory and pathogen transmission. PLoS Pathog 18, e1010939 (2022).

13. Koella, J. C., Sørensen, F. L. & Anderson, R. A. The malaria parasite, Plasmodium falciparum, increases the frequency of multiple feeding of its mosquito vector, Anopheles gambiae. Proc. R. Soc. Lond. B 265, 763–768 (1998).

14. Markwalter, C. F. et al. Plasmodium falciparum infection in humans and mosquitoes influence natural Anopheline biting behavior and transmission. Nat Commun 15, 4626 (2024).

15. Cator, L. J. et al. ‘Manipulation’ without the parasite: altered feeding behaviour of mosquitoes is not dependent on infection with malaria parasites. Proc. R. Soc. B. 280, 20130711 (2013).

16. Smallegange, R. C., et al. Malaria Infected Mosquitoes Express Enhanced Attraction to Human Odor. PLoS ONE 8, e63602 (2013).

17. Wekesa, J. W., Mwangi, R. W. & Copeland, R. S. Effect of Plasmodium Falciparum on Blood Feeding Behavior of Naturally Infected Anopheles Mosquitoes in Western Kenya. The American Journal of Tropical Medicine and Hygiene 47, 484–488 (1992).

18. Anderson, R. A., Koellaf, J. C. & Hurd, H. The effect of Plasmodium yoelii nigeriensis infection on the feeding persistence of Anopheles stephensi Liston throughout the sporogonic cycle. Proc. R. Soc. Lond. B 266, 1729–1733 (1999).

19. Ferguson, H. M. & Read, A. F. Mosquito appetite for blood is stimulated by Plasmodium chabaudi infections in themselves and their vertebrate hosts. Malaria Journal (2004).

20. Stanczyk, N. M. et al. Species-specific alterations in Anopheles mosquito olfactory responses caused by Plasmodium infection. Sci Rep 9, 3396 (2019).

21. Vantaux, A. et al. Host-seeking behaviors of mosquitoes experimentally infected with sympatric field isolates of the human malaria parasite Plasmodium falciparum: no evidence for host manipulation. Front. Ecol. Evol. 3, (2015).

22. Nguyen, P. L. et al. No evidence for manipulation of Anopheles gambiae, An. coluzzii and An. arabiensis host preference by Plasmodium falciparum. Sci Rep 7, 9415 (2017).

23. Vogels, C. B. et al. Virus interferes with host-seeking behaviour of mosquito. Journal of Experimental Biology 220, 3598–3603 (2017).

24. Yang, F., Chan, K., Brewster, C. C. & Paulson, S. L. Effects of La Crosse virus infection on the host-seeking behavior and levels of two neurotransmitters in Aedes triseriatus. Parasites & vectors 12, 1–7 (2019).

25. Putnam, J. L. & Scott, T. W. Blood-feeding behavior of dengue-2 virus-infected Aedes aegypti. The American journal of tropical medicine and hygiene 52, 225–227 (1995).

26. Platt, K. B. et al. Impact of dengue virus infection on feeding behavior of Aedes aegypti. The American journal of tropical medicine and hygiene 57, 119–125 (1997).

27. Sim, S., Ramirez, J. L. & Dimopoulos, G. Dengue virus infection of the Aedes aegypti salivary gland and chemosensory apparatus induces genes that modulate infection and blood-feeding behavior. PLoS pathogens 8, e1002631 (2012).

28. Maciel-de-Freitas, R., Sylvestre, G., Gandini, M. & Koella, J. C. The influence of dengue virus serotype-2 infection on Aedes aegypti (Diptera: Culicidae) motivation and avidity to blood feed. PLoS One 8, e65252 (2013).

29. Tallon, A. K. et al. Dengue infection modulates locomotion and host seeking in Aedes aegypti. PLoS neglected tropical diseases 14, e0008531 (2020).

30. Maire, T., Lambrechts, L. & Hol, F. J. Arbovirus impact on mosquito behavior: the jury is still out. Trends in Parasitology (2024).

31. Aleshnick, M., Ganusov, V. V., Nasir, G., Yenokyan, G. & Sinnis, P. Experimental determination of the force of malaria infection reveals a non-linear relationship to mosquito sporozoite loads. PLoS Pathog 16, e1008181 (2020).

32. Wei Xiang, B. W., et al. Dengue virus infection modifies mosquito blood-feeding behavior to increase transmission to the host. Proc. Natl. Acad. Sci. U.S.A. 119, e2117589119 (2022).

33. Hol, F. J., Lambrechts, L. & Prakash, M. BiteOscope, an open platform to study mosquito biting behavior. eLife 9, (2020).

34. Murray, G. P. D., Giraud, E. & Hol, F. J. H. Characterizing Mosquito Biting Behavior Using the BiteOscope. Cold Spring Harb Protoc 2023, pdb.prot108176 (2023).

35. Maire, T., Wan, Z., Lambrechts, L. & Hol, F. J. BuzzWatch: Uncovering Multi-scale Temporal Patterns in Mosquito Behavior Through Continuous Long-term Monitoring. Preprint at 10.7554/eLife.107916.1 (2025).

36. Mathis, A. et al. DeepLabCut: markerless pose estimation of user-defined body parts with deep learning. Nat Neurosci 21, 1281–1289 (2018).

37. Pereira, T. D. et al. SLEAP: A deep learning system for multi-animal pose tracking. Nature methods 19, 486–495 (2022).

38. Baik, L. S. et al. Mosquito taste responses to human and floral cues guide biting and feeding. Nature 635, 639–646 (2024).

39. Smith, D. L. & Ellis McKenzie, F. Statics and dynamics of malaria infection in Anopheles mosquitoes. Malar J 3, 13 (2004).

40. Tjaden, N. B., Thomas, S. M., Fischer, D. & Beierkuhnlein, C. Extrinsic Incubation Period of Dengue: Knowledge, Backlog, and Applications of Temperature Dependence. PLoS Negl Trop Dis 7, e2207 (2013).

41. Ohm, J. R. et al. Rethinking the extrinsic incubation period of malaria parasites. Parasites Vectors 11, 178 (2018).

42. Graumans, W., Jacobs, E., Bousema, T. & Sinnis, P. When Is a Plasmodium-Infected Mosquito an Infectious Mosquito? Trends in Parasitology 36, 705–716 (2020).

43. Sakuma, C. et al. Fibrinopeptide A-induced blood-feeding arrest in the yellow fever mosquito Aedes aegypti. Cell Reports 43, 114354 (2024).

44. Keeling, M. J. & Rohani, P. Modeling Infectious Diseases in Humans and Animals. (Princeton university press, 2008).

45. Mandal, S., Sarkar, R. R. & Sinha, S. Mathematical models of malaria-a review. Malaria journal 10, 1–19 (2011).

46. Sinka, M. E., et al. A new malaria vector in Africa: Predicting the expansion range of Anopheles stephensi and identifying the urban populations at risk. Proc. Natl. Acad. Sci. U.S.A. 117, 24900–24908 (2020).

47. Emiru, T. et al. Evidence for a role of Anopheles stephensi in the spread of drug- and diagnosis-resistant malaria in Africa. Nat Med 29, 3203–3211 (2023).

48. Ponnudurai, T., Meuwissen, J. T., Leeuwenberg, A. D., Verhave, J. & Lensen, A. The production of mature gametocytes of Plasmodium falciparum in continuous cultures of different isolates infective to mosquitoes. Transactions of the Royal Society of Tropical Medicine and Hygiene 76, 242–250 (1982).

49. Ponnudurai, T. et al. Infectivity of cultured Plasmodium falciparum gametocytes to mosquitoes. Parasitology 98, 165–173 (1989).

50. Teirlinck, A. C. et al. NF135.C10: A New Plasmodium falciparum Clone for Controlled Human Malaria Infections. The Journal of Infectious Diseases 207, 656–660 (2013).

51. Van De Vegte-Bolmer, M. et al. A portfolio of geographically distinct laboratory-adapted Plasmodium falciparum clones with consistent infection rates in Anopheles mosquitoes. Malar J 20, 381 (2021).

52. Minassian, A. M. et al. Controlled human malaria infection with a clone of Plasmodium vivax with high-quality genome assembly. JCI Insight 6, e152465 (2021).

53. Graumans, W. et al. Semi-high-throughput detection of Plasmodium falciparum and Plasmodium vivax oocysts in mosquitoes using bead-beating followed by circumsporozoite ELISA and quantitative PCR. Malar J 16, 356 (2017).

54. Hermsen, C. C. et al. Detection of Plasmodium falciparum malaria parasites in vivo by real-time quantitative PCR. Molecular and Biochemical Parasitology 118, 247–251 (2001).

55. Wampfler, R. et al. Strategies for Detection of Plasmodium species Gametocytes. PLoS ONE 8, e76316 (2013).

56. Fontaine, A., Jiolle, D., Moltini-Conclois, I., Lequime, S. & Lambrechts, L. Excretion of dengue virus RNA by Aedes aegypti allows non-destructive monitoring of viral dissemination in individual mosquitoes. Sci Rep 6, 24885 (2016).

57. De La Llave, E. et al. A combined luciferase imaging and reverse transcription polymerase chain reaction assay for the study of Leishmania amastigote burden and correlated mouse tissue transcript fluctuations: A combined bioluminescence/RT-PCR assay. Cellular Microbiology 13, 81–91 (2011).

58. Giraud, E., Martin, O., Yakob, L. & Rogers, M. Quantifying Leishmania Metacyclic Promastigotes from Individual Sandfly Bites Reveals the Efficiency of Vector Transmission. Commun Biol 2, 84 (2019).

59. Gentile, C., Lima, J. & Peixoto, A. Isolation of a fragment homologous to the rp49 constitutive gene of Drosophila in the Neotropical malaria vector Anopheles aquasalis (Diptera: Culicidae). Mem. Inst. Oswaldo Cruz 100, 545–547 (2005).

60. Merkling, S. H. et al. Dengue virus susceptibility in Aedes aegypti linked to natural cytochrome P450 promoter variants. Nat Commun 16, 7468 (2025).

61. Jolles, J. pirecorder: Controlled and automated image and video recording with the raspberry pi. JOSS 5, 2584 (2020).

62. Lauer, J. et al. Multi-animal pose estimation, identification and tracking with DeepLabCut. Nat Methods 19, 496–504 (2022).

63. Simonyan, K., Vedaldi, A. & Zisserman, A. Deep Inside Convolutional Networks: Visualising Image Classification Models and Saliency Maps. Preprint at 10.48550/ARXIV.1312.6034 (2013).

64. Doolan, D. L., Dobaño, C. & Baird, J. K. Acquired Immunity to Malaria. Clin Microbiol Rev 22, 13–36 (2009).

65. Ghani, A. C. et al. Loss of Population Levels of Immunity to Malaria as a Result of Exposure-Reducing Interventions: Consequences for Interpretation of Disease Trends. PLoS ONE 4, e4383 (2009).

66. Soetaert, K., Petzoldt, T. & Setzer, R. W. Solving Differential Equations in *R* : Package **deSolve**. J. Stat. Soft. 33, (2010).

67. Obadia, T., Haneef, R. & Boëlle, P.-Y. The R0 package: a toolbox to estimate reproduction numbers for epidemic outbreaks. BMC Med Inform Decis Mak 12, 147 (2012).

68. Cori, A., Ferguson, N. M., Fraser, C. & Cauchemez, S. A New Framework and Software to Estimate Time-Varying Reproduction Numbers During Epidemics. American Journal of Epidemiology 178, 1505–1512 (2013).

69. Andolina, C. et al. Quantification of sporozoite expelling by Anopheles mosquitoes infected with laboratory and naturally circulating P. falciparum gametocytes. eLife 12, RP90989 (2024).

70. Putnam, J. L. & Scott, T. W. The Effect of Multiple Host Contacts on the Infectivity of Dengue-2 Virus-Infected Aedes aegypti. The Journal of Parasitology 81, 170 (1995).

71. Feldmann, A. M. & Ponnudurai, T. Selection of Anopheles stephensi for refractoriness and susceptibility to Plasmodium falciparum. Medical Vet Entomology 3, 41–52 (1989).

72. Alout, H. et al. Interactive cost of Plasmodium infection and insecticide resistance in the malaria vector Anopheles gambiae. Sci Rep 6, 29755 (2016).

73. Meister, S. et al. Anopheles gambiae PGRPLC-Mediated Defense against Bacteria Modulates Infections with Malaria Parasites. PLoS Pathog 5, e1000542 (2009).

74. Lambrechts, L. et al. Genetic specificity and potential for local adaptation between dengue viruses and mosquito vectors. BMC Evol Biol 9, 160 (2009).

75. Fansiri, T. et al. Genetic Mapping of Specific Interactions between Aedes aegypti Mosquitoes and Dengue Viruses. PLoS Genet 9, e1003621 (2013).

76. Anderson, R. M. & May, R. M. Infectious Diseases of Humans: Dynamics and Control. (Oxford University PressOxford, 1991). doi:10.1093/oso/9780198545996.001.0001.

77. Pandey, A., Mubayi, A. & Medlock, J. Comparing vector–host and SIR models for dengue transmission. Mathematical Biosciences 246, 252–259 (2013).

78. Chitnis, N., Hyman, J. M. & Cushing, J. M. Determining Important Parameters in the Spread of Malaria Through the Sensitivity Analysis of a Mathematical Model. Bull. Math. Biol. 70, 1272– 1296 (2008).

79. Filipe, J. A. N., Riley, E. M., Drakeley, C. J., Sutherland, C. J. & Ghani, A. C. Determination of the Processes Driving the Acquisition of Immunity to Malaria Using a Mathematical Transmission Model. PLoS Comput Biol 3, e255 (2007).

80. Cummins, B., Cortez, R., Foppa, I. M., Walbeck, J. & Hyman, J. M. A Spatial Model of Mosquito Host-Seeking Behavior. PLoS Comput Biol 8, e1002500 (2012).

