## Supplementary Information for "Deep behavioral phenotyping of pathogen infected mosquitoes reveals species-specific behavior changes enhancing transmission"

### Supplementary File

**Figure S1. Survival mosquito overtime in BuzzWatch setup.**

Total number of survival mosquitoes for uninfected (Ctrl) and infected (Infection) cages in BuzzWatch setup were recorded 4 times throughout the 3-week long BuzzWatch experiments, namely at 0 dpi, 7 dpi, 14 dpi and 21 dpi. Recording was done by visual inspection when replacing the sugar feeder.

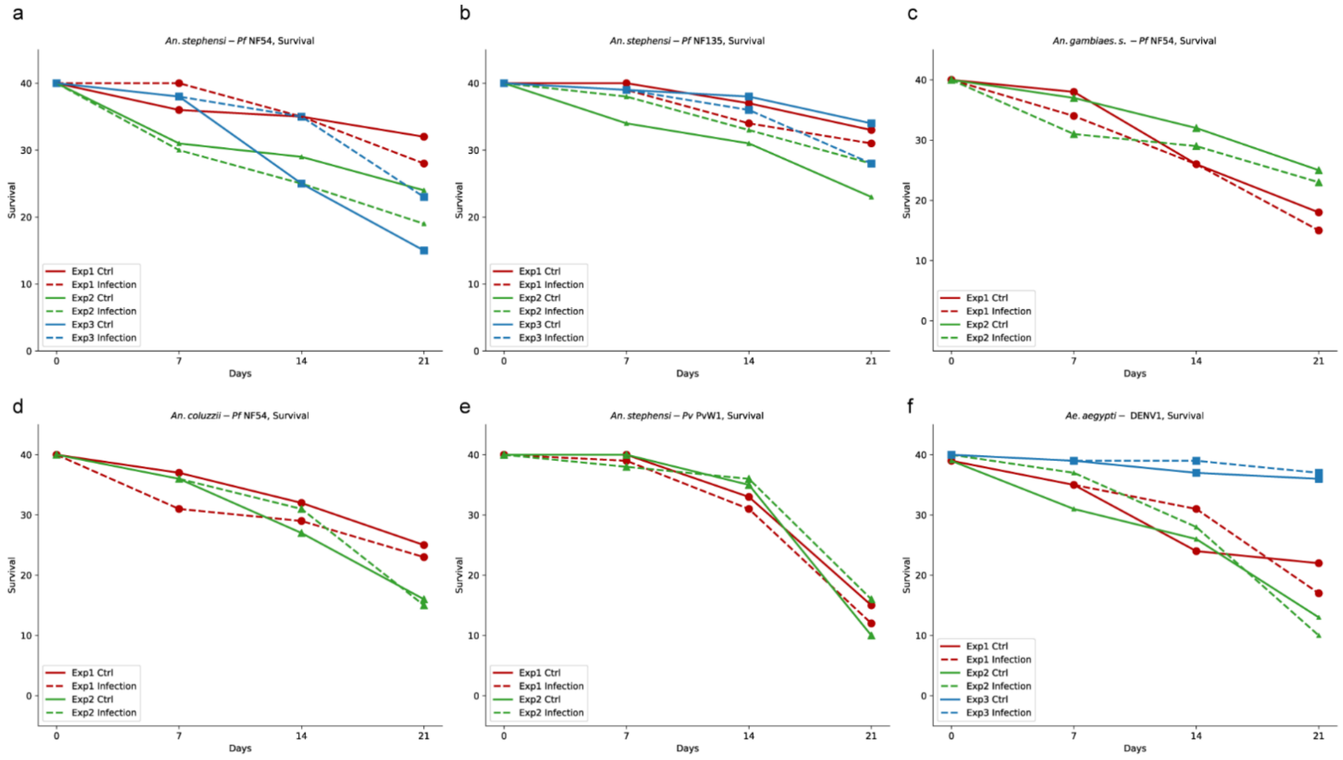

**Figure S2. Fraction of flying mosquitoes over the long-term monitoring (Buzzwatch) experiment**

Averaged activity curves of all 21 days for uninfected (Ctrl) and infected (Infection) *An. stephensi* – *Pf* NF135 (a), *An. gambiae s.s.* – *Pf* NF54 (b), *An. coluzzii.* – *Pf* NF54 (c), 2 replicates of *An. stephensi* – *Pv* PvW1 (d,e) and *Ae. aegypti* – DENV1 (f). We report the 2 replicates of *An. stephensi* – *Pv* PvW1 separately as these two experiments were conducted in different climate room with slightly different dimming transitions in the light cycles.

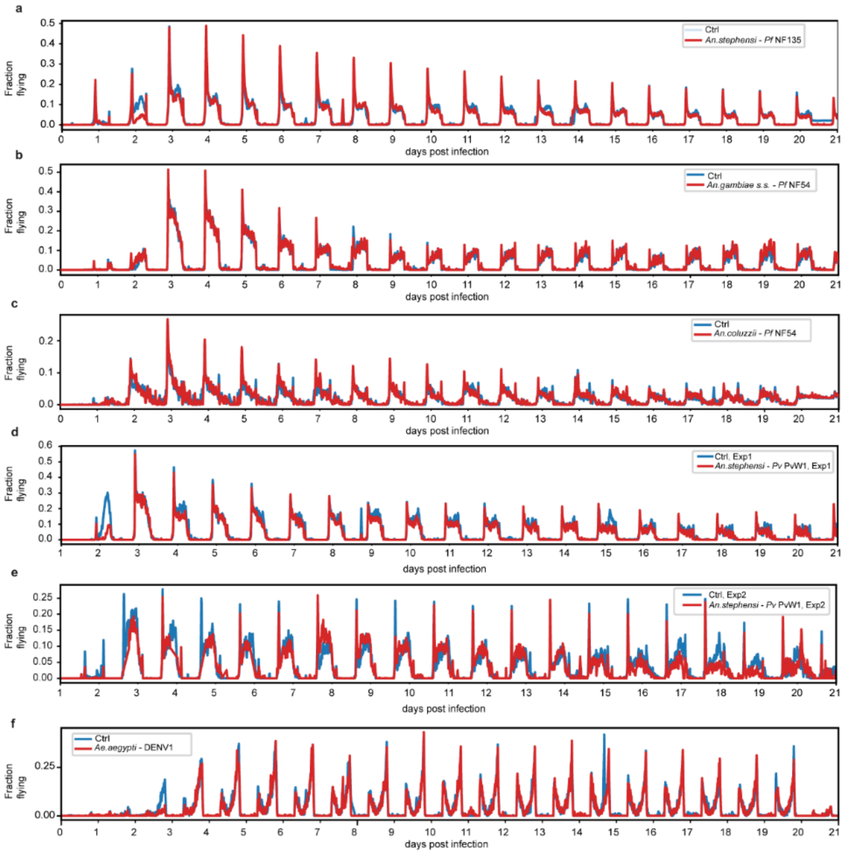

**Figure S3. Daily average of fraction of flying mosquito in BuzzWatch experiment**

Mean flight activity per day for uninfected (Ctrl) and infected (Infection) *An. stephensi – Pf* NF135 (a), *An. gambiae s.s. – Pf* NF54 (b), *An. coluzzii. – Pf* NF54 (c), 2 replicates of *An. stephensi – Pv* PvW1 (d,e) and *Ae. aegypti* – DENV1 (f). 2 replicates of *An. stephensi – Pv* PvW1 were reported separately with Zeitgeber time adjusted according to the climate room day-night light cycle.

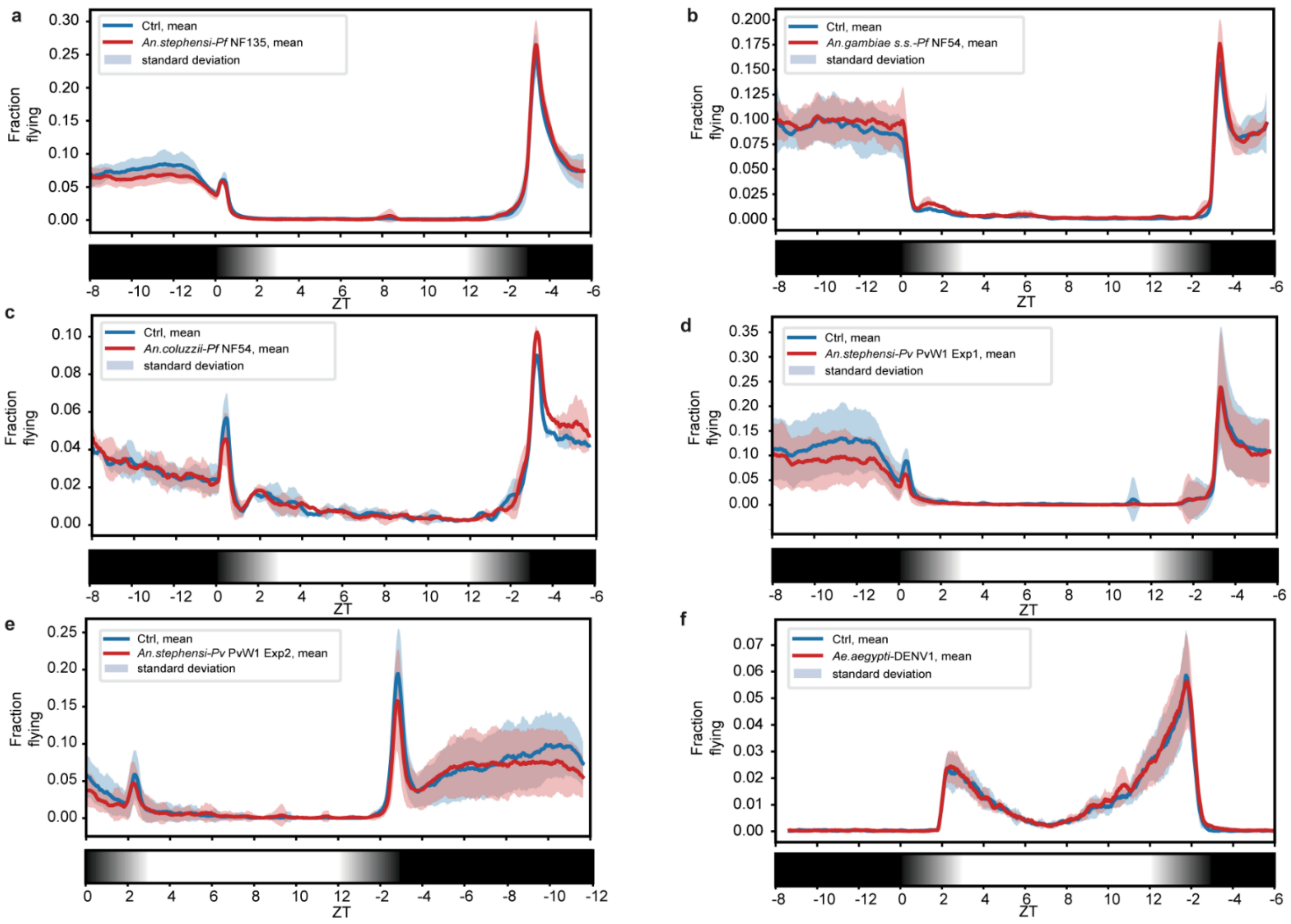

**Figure S4. Daily average of sugar feeding index in BuzzWatch experiment**

Daily average of the sugar feeding index for uninfected (Ctrl) and infected (Infection) *An. stephensi* – *Pf* NF135 (a), *An. gambiae s.s.* – *Pf* NF54 (b), *An. coluzzii.* – *Pf* NF54 (c), 2 replicates of *An. stephensi* – *Pv* PvW1 (d,e) and *Ae. aegypti* – DENV1 (f). 2 replicates of *An. stephensi* – *Pv* PvW1 were reported accordingly as indicated in Fig S3 and S4.

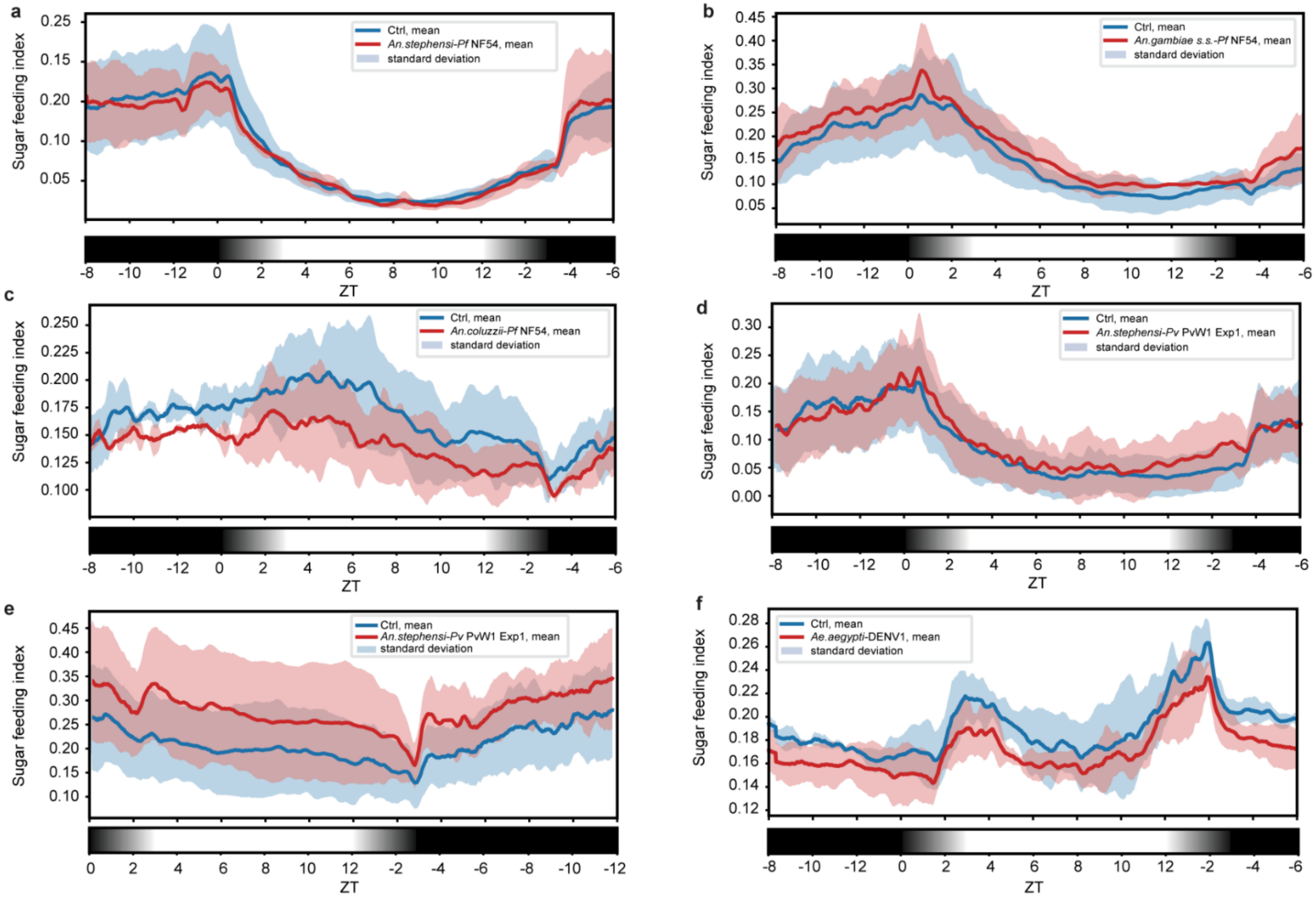

**Figure S5. Z-score BuzzWatch – z-score table**

Heatmap of GLMM test for 6 different quantitative metrics characterizing mosquito flight and sugar feeding behavior of uninfected (Ctrl) and infected (Infection) *An. stephensi – Pf* NF135 (a), *An. gambiae s.s. – Pf* NF54 (b), *An. coluzzii. – Pf* NF54 (c) and *Ae. aegypti* – DENV1 (f). Due to the heterogeneity in zeigeber time, we did not perform GLMM test for 2 replicates of *An. stephensi – Pv* PvW1.

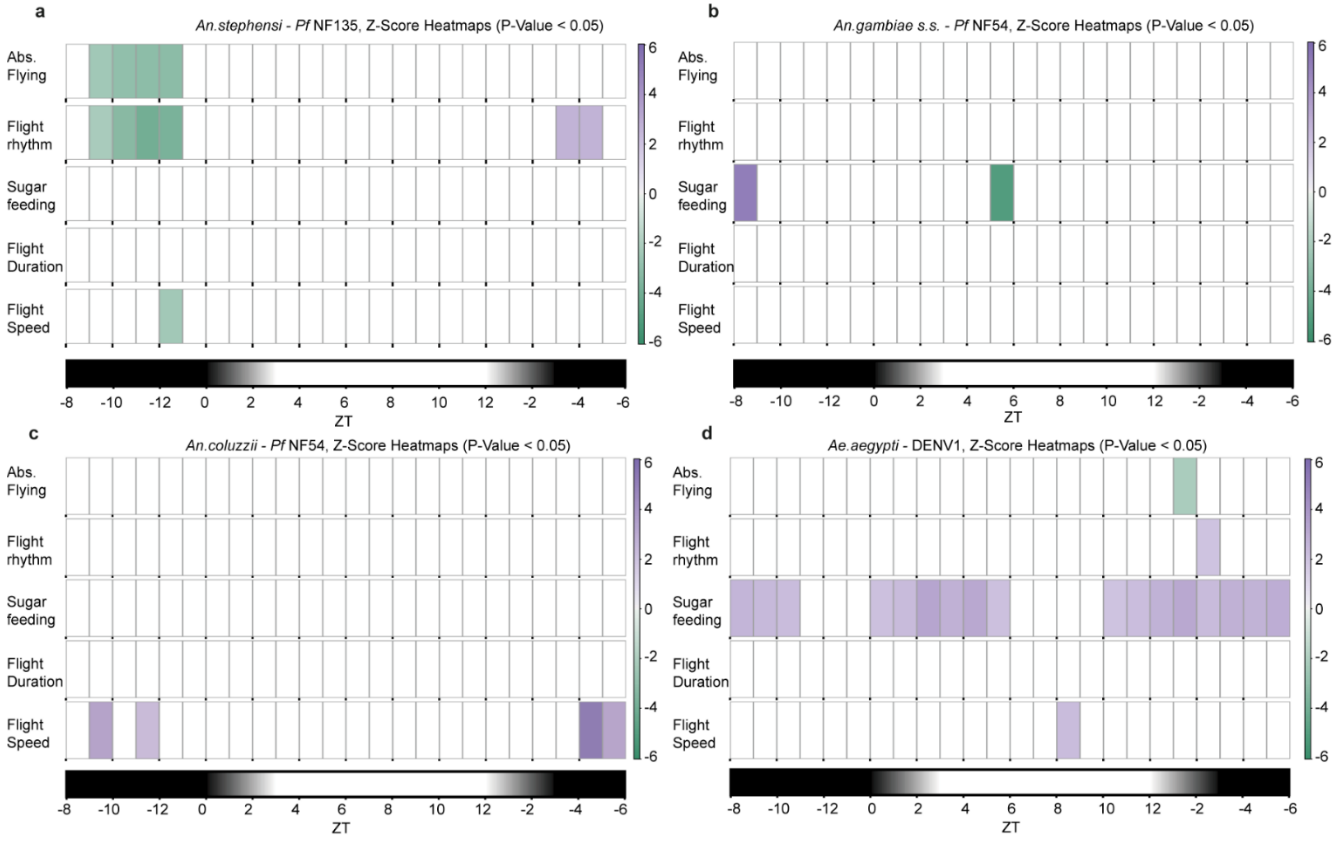

**Figure S6. Walking time, probing time and engorge time of engorged individual mosquitoes in BiteOscope experiment**

Behavioral decomposition of tracks leading to engorgement into walking, probing, and engorging time with different colors representing distinct behaviors of uninfected (Ctrl) and infected (Infection)*An. stephensi – Pf* NF135 (a), *An. gambiae s.s. – Pf* NF54 (b), *An. coluzzii. – Pf* NF54 (c), *An. stephensi – Pv* PvW1 (d) and *Ae. aegypti* – DENV1 (e). No significant differences on these parameters were observed on these mosquito-pathogen combinations.

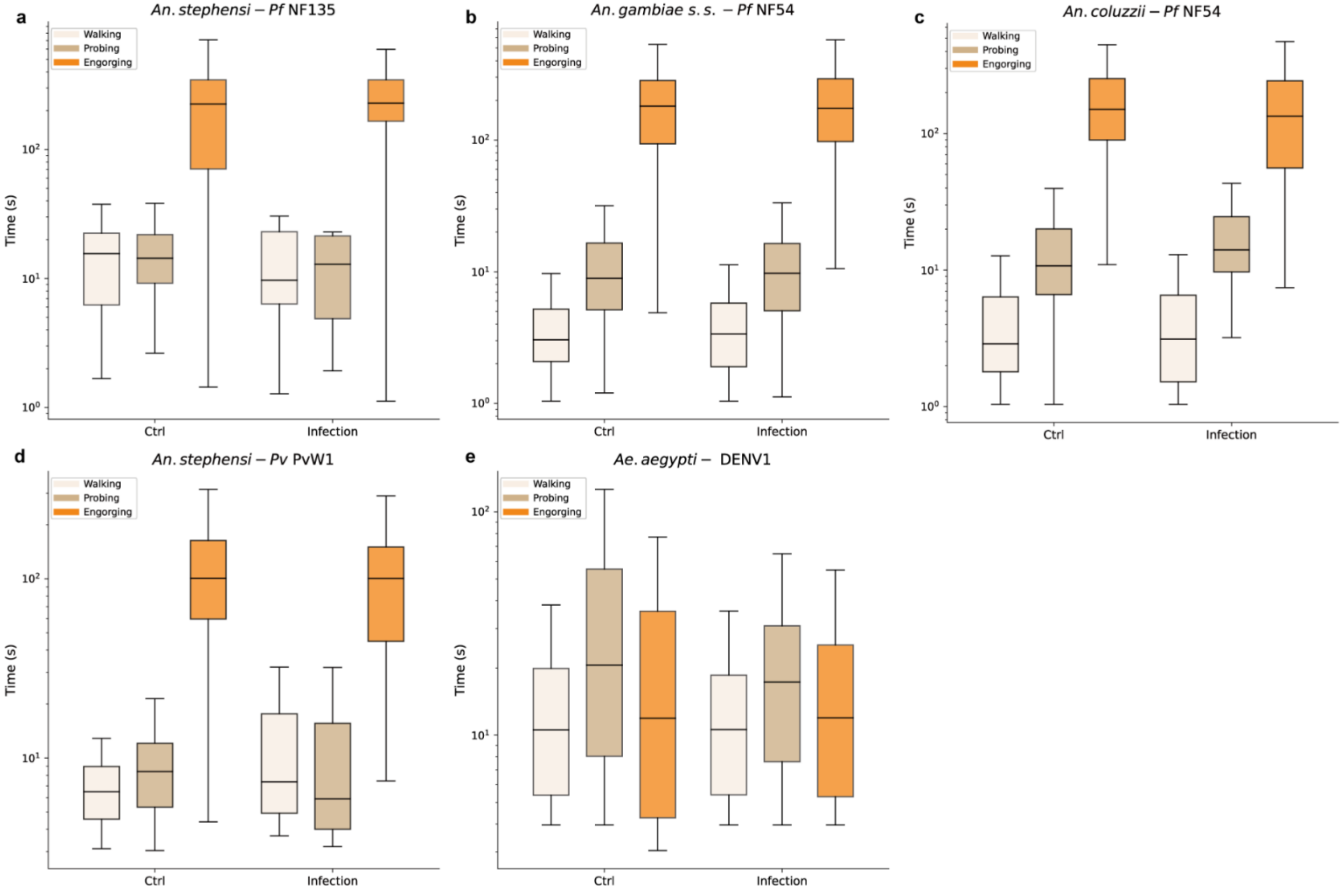

**Figure S7. Mean speed and moving fraction of engorged individual mosquitoes in BiteOscope experiment**

Mean speed and moving fraction of engorged mosquitoes of uninfected (Ctrl) and infected (Infection)*An. stephensi – Pf* NF135 (a), *An. gambiae s.s. – Pf* NF54 (b), *An. coluzzii. – Pf* NF54 (c), *An. stephensi – Pv* PvW1 (d) and *Ae. aegypti* – DENV1 (e). No significant differences on these parameters were observed on these mosquito-pathogen combinations.

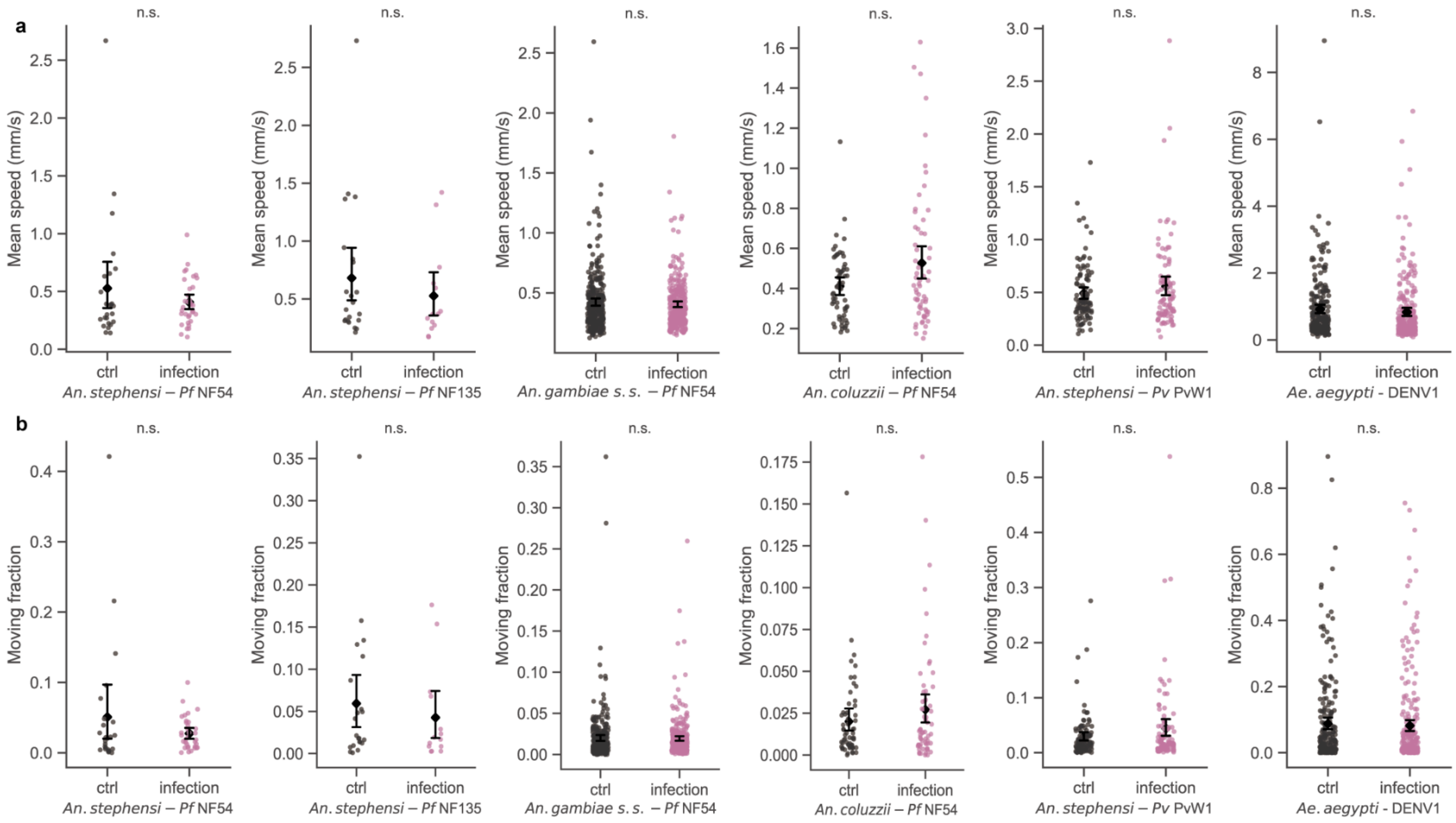

**Figure S8. Maximum abdomen width and abdomen increase of engorged individual mosquitoes in BiteOscope experiment**

Maximum abdomen width and abdomen increase of engorged mosquitoes of uninfected (Ctrl) and infected (Infection)*An. stephensi – Pf* NF135 (a), *An. gambiae s.s. – Pf* NF54 (b), *An. coluzzii. – Pf* NF54 (c), *An. stephensi – Pv* PvW1 (d) and *Ae. aegypti* – DENV1 (e). No significant differences on these parameters were observed on these mosquito-pathogen combinations.

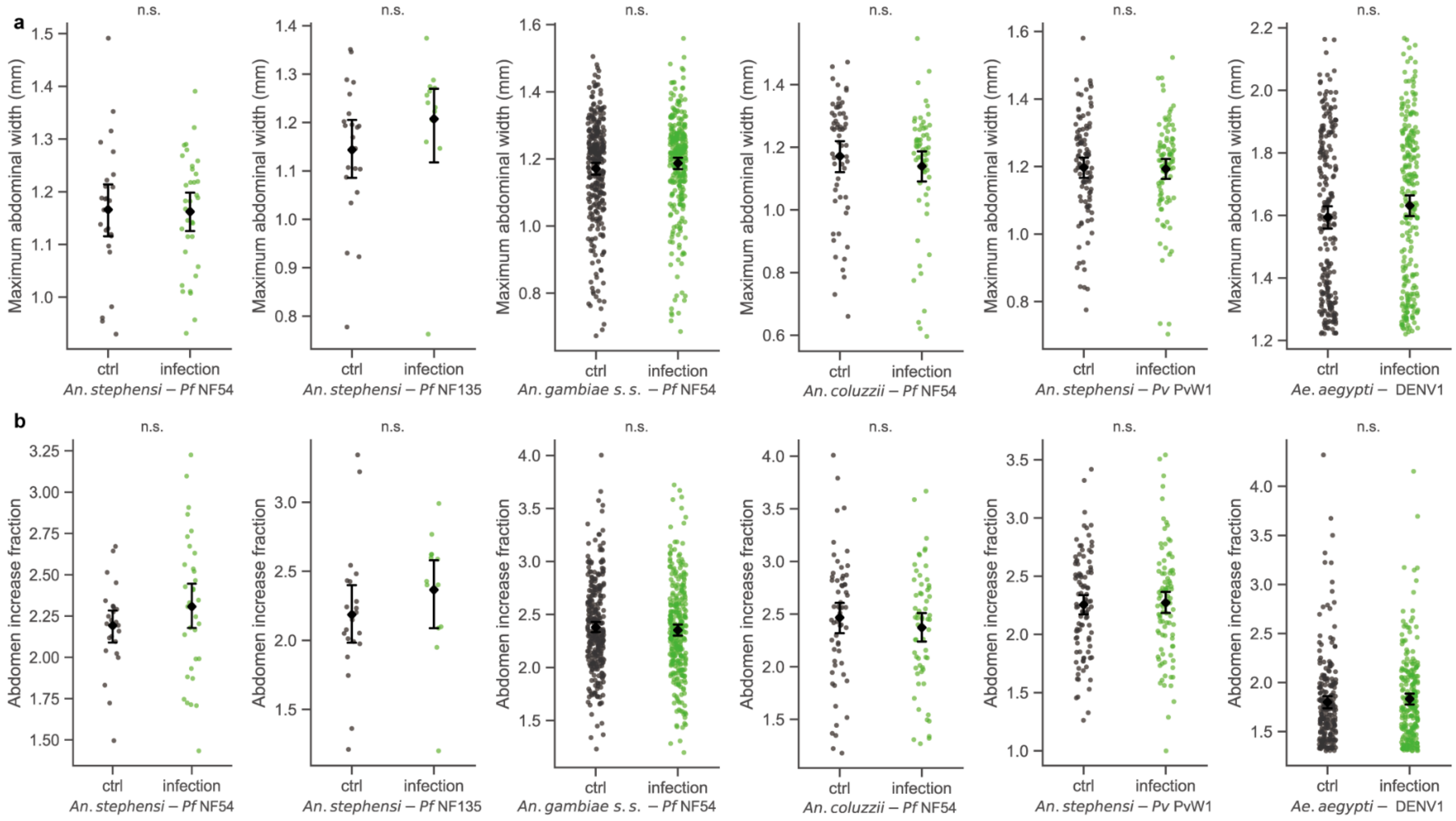

**Figure S9. Total time, total distance, mean speed and moving fraction of no-engorge individual mosquitoes in BiteOscope experiment**

Total time, total distance, mean speed and moving fraction of no-engorge mosquitoes of uninfected (Ctrl) and infected (Infection)*An. stephensi* – *Pf* NF135 (a), *An. gambiae s.s.* – *Pf* NF54 (b), *An. coluzzii.* – *Pf* NF54 (c), *An. stephensi* – *Pv* PvW1 (d) and *Ae. aegypti* – DENV1 (e). No significant differences on these parameters were observed on these mosquito-pathogen combinations.

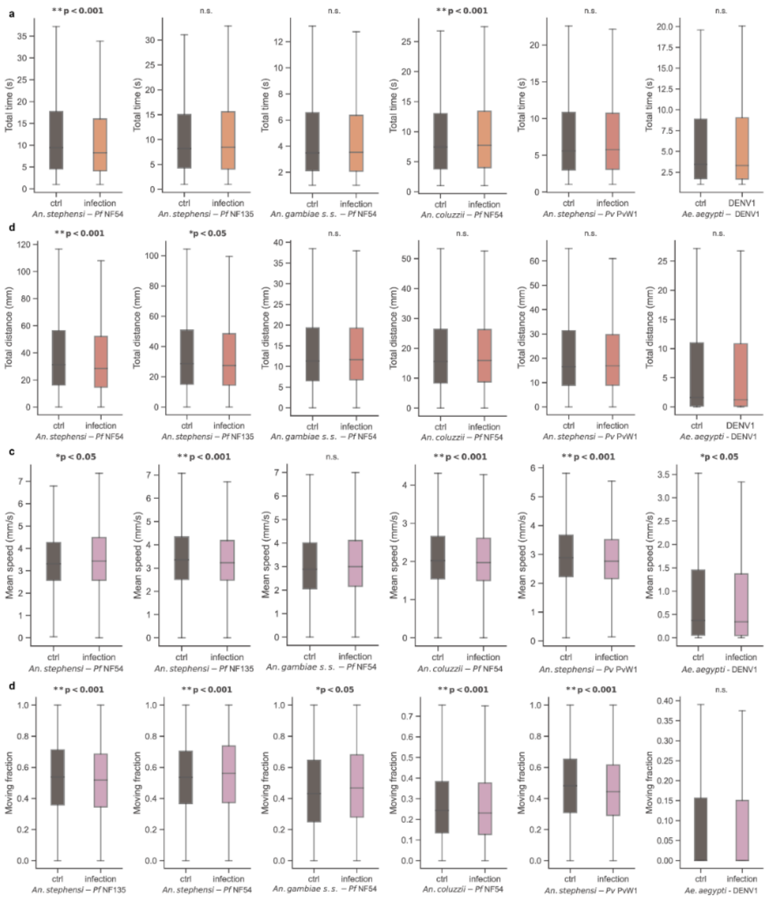

**Figure S10. Temporal dynamics of infected (I) human populations under different mosquito biting rates in the mathematical model of malaria and dengue fever.**

Modelling of temporal dynamics of infected individual number in the human population under scenarios incorporating Anopheles mosquito behavioral alterations (a) and Aedes mosquito behavioral alterations (b).

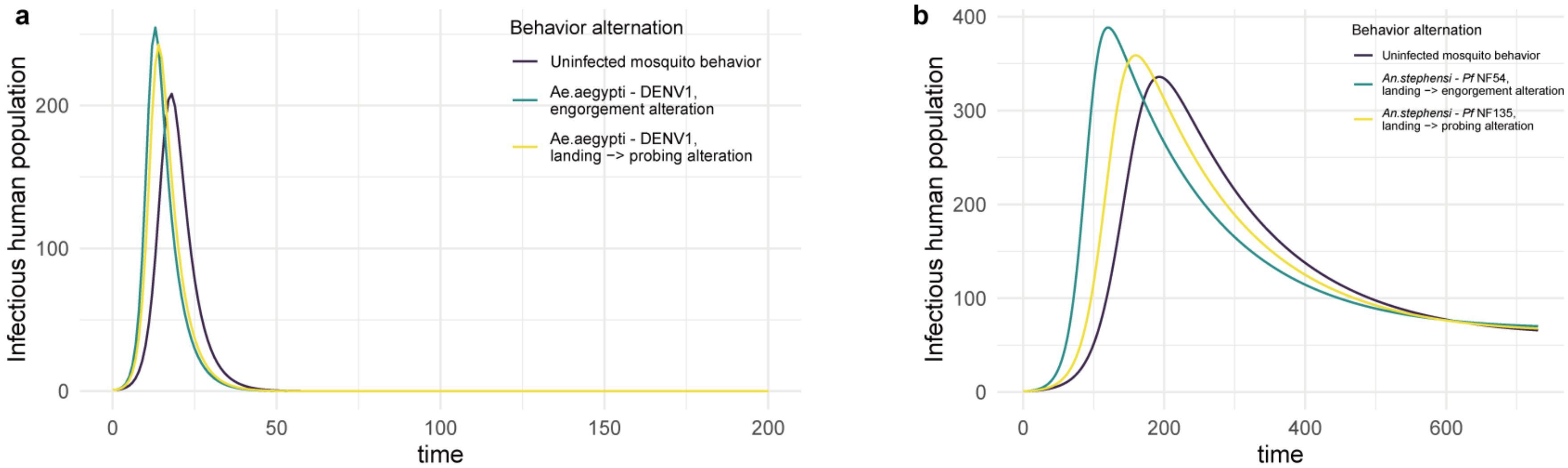

**Table S1. Key resources material**

| Animal species, reagents and computational tool | Source or reference |
|---|---|
| Organisms/strains | |
| *Anopheles stephensi* (Sind-Kasur) | Feldmann and Ponnudurai, 1989 [71] |
| *Anopheles gambiae s.s.* (Kisumu) | Alout et al. 2016 [72] |
| *Anopheles coluzzii* (Ngousso) | Meister et al. 2009 [73] |
| *Aedes aegypti* (KPP-Thailand TN) | Lambrechts et al. 2009 [74] |
| *Plasmodium falciparum* (Nijmegen falciparum 54) | Ponnudurai et al. 1982 [48]<br>Ponnudurai et al. 1989 [49] |
| *Plasmodium falciparum* (Nijmegen falciparum 135) | Teirlinck et al. 2013 [50] |
| *Plasmodium vivax* (PvW1) | Minassian et al. 2021 [52] |
| *Orthoflavivirus dengue* - Dengue virus (DENV-1 isolate KDH0026A) | Fansiri et al. 2013 [75] |
| Software/Data analysis | |
| BuzzWatch codes | Maire et al. 2025 [35]<br>Available from: https://theomaire.github.io/buzzwatch/construct.html |
| BiteOscope codes | Available from: https://github.com/felixhol/biteOscope |
| BiteOServer codes | Available from: https://github.com/Vlad-2299/bite-o-serve |
| Deeplabcut | Mathis et al. 2018, Lauer et al. 2022. [36,62]<br>Version 2.3.5, Available from: https://github.com/deeplabcut |

**Table S2. BiteOscope experiments outcomes and populational-level behavioral statistics.**

| | *An. stephensi* - Pf NF54 | | *An. stephensi* - Pf NF135 | | *An. gambiae* s.s. - Pf NF54 | | An.coluzzii - Pf NF54 | | *An. stephensi* - *Pv* PvW1 | | *Aedes aegypti* - DENV | |
|---|---|---|---|---|---|---|---|---|---|---|---|---|
| | Uninfected | Infected | Uninfected | Infected | Uninfected | Infected | Uninfected | Infected | Uninfected | Infected | Uninfected | Infected |
| N | 350 | 350 | 350 | 350 | 550 | 550 | 421 | 414 | 250 | 250 | 759 | 519 |
| N tracks (Landing) | 5127 | 3815 | 5156 | 4797 | 12298 | 11881 | 19274 | 20887 | 5717 | 5390 | 1949 | 2157 |
| N behavior classification tracks | 3330 | 2299 | 3097 | 2844 | 2219 | 2229 | 7848 | 7563 | 2290 | 1476 | 890 | 849 |
| N engorgement tracks | 24 | 34 | 23 | 15 | 312 | 297 | 56 | 62 | 100 | 92 | 223 | 220 |
| N probing tracks (No engorgement) | 465 | 323 | 389 | 446 | 648 | 704 | 5294 | 5167 | 409 | 254 | 198 | 218 |
| landing rate (per mosquito) | 11 | 15 | 14 | 15 | 22 | 22 | 46 | 50 | 22 | 23 | 2.7 | 4.3 |
| | $p = 0.12$ | | $p = 0.85$ | | $p = 0.71$ | | $p = 0.28$ | | p = 0.88 | | $p < 0.05$* | |
| engorgement ratio | $p = 0.22$ | | $p = 0.24$ | | $p = 0.40$ | | $p = 0.55$ | | p = 0.52 | | $p < 0.001$** | |
| | OR = 1.46 | | OR = 0.63 | | OR = 0.90 | | OR = 1.15 | | OR = 0.87 | | OR = 1.77 | |
| landing (no engorgement) → probing (no engorgement) | $p = 0.84$ | | $p < 0.001$** | | $p = 1.11$ | | $p = 0.21$ | | $p = 0.83$ | | $p < 0.05$* | |
| | OR = 1.02 | | OR = 1.29 | | OR = 0.11 | | OR = 1.04 | | OR = 0.98 | | OR = 1.52 | |
| landing → engorgement | $p < 0.05$* | | $p = 0.33$ | | $p = 0.87$ | | $p = 0.36$ | | $p = 0.88$ | | $p = 0.21$ | |
| | OR = 1.92 | | OR = 0.70 | | OR = 0.99 | | OR = 1.20 | | OR = 0.97 | | OR = 0.87 | |

**Table S3. Individual-level behavior statistics of all mosquitoes successfully engorged.**

| | | *Anopheles stephensi* - Pf NF54 | | | *Anopheles stephensi* - Pf NF135 | | | *Anopheles gambiae s.s.* - Pf NF54 | | | *Anopheles coluzzii* - Pf NF54 | | | *Anopheles stephensi* - Pv PvW1 | | | *Aedes aegypti* - DENV | | |
|---|---|---|---|---|---|---|---|---|---|---|---|---|---|---|---|---|---|---|---|
| | | Mean | std | p | Mean | std | p | Mean | std | p | Mean | std | p | Mean | std | p | Mean | std | p |
| total time (s) | Uninfected | 355.08 | 193.68 | < 0.05* | 363.76 | 244.67 | 0.65 | 305.41 | 204.89 | 0.66 | 274.31 | 166.14 | 0.78 | 272.70 | 139.24 | 0.58 | 63.90 | 82.94 | 0.43 |
| | Infected | 509.91 | 265.62 | | 399.68 | 270.13 | | 307.37 | 218.71 | | 269.24 | 178.82 | | 267.68 | 152.09 | | 55.54 | 48.20 | |
| walking time (s) | Uninfected | 12.87 | 14.41 | 0.89 | 17.07 | 13.11 | 0.61 | 5.66 | 21.56 | 0.67 | 17.07 | 13.11 | 0.61 | 9.55 | 10.19 | 0.29 | 21.24 | 44.48 | 0.82 |
| | Infected | 10.07 | 10.10 | | 16.43 | 15.27 | | 5.97 | 16.06 | | 16.43 | 15.27 | | 14.24 | 14.42 | | 17.19 | 24.53 | |
| probing time (s) | Uninfected | 7.94 | 6.36 | 0.35 | 16.64 | 9.55 | 0.58 | 13.22 | 13.07 | 0.71 | 16.64 | 9.55 | 0.58 | 10.11 | 6.80 | 0.48 | 38.35 | 43.12 | 0.15 |
| | Infected | 11.41 | 11.46 | | 20.83 | 22.27 | | 13.05 | 11.61 | | 20.83 | 22.27 | | 11.52 | 10.33 | | 31.33 | 48.98 | |
| Average probing bouts per landing | Uninfected | 1.80 | 1.23 | 0.09 | 3.72 | 2.24 | 0.90 | 3.22 | 3.27 | 0.92 | 2.82 | 2.11 | 0.05 | 3.62 | 2.66 | 0.87 | 4.87 | 7.15 | 0.85 |
| | Infected | 2.65 | 1.94 | | 4.21 | 3.34 | | 2.44 | 2.66 | | 3.58 | 2.43 | | 3.80 | 3.07 | | 4.62 | 6.46 | |
| engorging time (s) | Uninfected | 168.45 | 108.65 | < 0.05* | 238.12 | 188.92 | 0.56 | 216.63 | 165.17 | 0.99 | 238.12 | 188.92 | 0.56 | 121.59 | 84.36 | 0.32 | 37.87 | 77.78 | 0.71 |
| | Infected | 241.79 | 134.34 | | 288.47 | 214.11 | | 222.97 | 177.41 | | 288.47 | 214.11 | | 106.39 | 78.04 | | 25.72 | 39.36 | |
| total walking distance (mm) | Uninfected | 133.50 | 74.78 | < 0.05* | 163.32 | 72.90 | 0.65 | 107.92 | 67.85 | 0.87 | 100.18 | 55.66 | 0.42 | 117.52 | 66.89 | 0.78 | 33.81 | 35.40 | 0.32 |
| | Infected | 182.33 | 90.32 | | 162.31 | 96.27 | | 105.18 | 60.36 | | 114.24 | 73.81 | | 128.13 | 83.92 | | 33.72 | 28.39 | |
| moving fraction | Uninfected | 0.05 | 0.09 | 0.65 | 0.06 | 0.08 | 0.56 | 0.02 | 0.03 | 0.58 | 0.02 | 0.02 | 0.45 | 0.03 | 0.04 | 0.34 | 0.08 | 0.14 | 0.25 |
| | Infected | 0.03 | 0.02 | | 0.04 | 0.05 | | 0.02 | 0.02 | | 0.03 | 0.03 | | 0.04 | 0.08 | | 0.09 | 0.14 | |

| | | | | | | | | | | | | | | | | | | | |
|---|---|---|---|---|---|---|---|---|---|---|---|---|---|---|---|---|---|---|---|
| mean speed (mm/s) | Uninfected | 0.53 | 0.53 | 0.09 | 0.69 | 0.59 | 0.43 | 0.43 | 0.27 | 0.96 | 0.41 | 0.17 | 0.23 | 0.49 | 0.28 | 0.69 | 0.84 | 0.95 | 0.07 |
| | Infected | 0.41 | 0.20 | | 0.53 | 0.38 | | 0.41 | 0.20 | | 0.53 | 0.34 | | 0.56 | 0.42 | | 0.93 | 1.03 | |
| maximum abdomen width (mm) | Uninfected | 1.17 | 0.13 | 0.87 | 1.14 | 0.14 | 0.06 | 1.17 | 0.16 | 0.26 | 1.17 | 0.19 | 0.13 | 1.20 | 0.15 | 0.88 | 1.63 | 0.26 | 0.16 |
| | Infected | 1.16 | 0.11 | | 1.21 | 0.14 | | 1.19 | 0.15 | | 1.14 | 0.19 | | 1.20 | 0.14 | | 1.59 | 0.26 | |
| abdomen increase | Uninfected | 2.19 | 0.26 | 0.22 | 2.19 | 0.47 | 0.09 | 2.35 | 0.44 | 0.52 | 2.47 | 0.60 | 0.35 | 2.26 | 0.43 | 0.92 | 1.83 | 0.44 | 0.25 |
| | Infected | 2.31 | 0.42 | | 2.37 | 0.44 | | 2.38 | 0.43 | | 2.37 | 0.55 | | 2.27 | 0.47 | | 1.80 | 0.45 | |

**Table S4. Individual-level behavior statistics of all mosquitoes without engorgement.**

| | | *Anopheles stephensi* - Pf NF54 | | | *Anopheles stephensi* - Pf NF135 | | | *Anopheles gambiae s.s.* - Pf NF54 | | | *Anopheles coluzzii* - Pf NF54 | | | *Anopheles stephensi* - *Pv* PvW1 | | | *Aedes aegypti* - DENV | | |
|---|---|---|---|---|---|---|---|---|---|---|---|---|---|---|---|---|---|---|---|
| | | Mean | std | p | Mean | std | p | Mean | std | p | Mean | std | p | Mean | std | p | Mean | std | p |
| total time (s) | Uninfected | 13.77 | 14.70 | < 0.001** | 11.72 | 11.75 | 0.98 | 5.46 | 7.93 | 0.95 | 9.77 | 8.75 | < 0.001** | 9.06 | 11.28 | 0.27 | 10.74 | 35.05 | 0.42 |
| | Infected | 12.56 | 13.54 | | 12.07 | 12.34 | | 5.56 | 7.83 | | 10.01 | 8.86 | | 8.87 | 10.37 | | 10.85 | 31.70 | |
| walking time (s) | Uninfected | 10.84 | 8.33 | 0.82 | 9.90 | 6.84 | < 0.05* | 6.19 | 11.07 | 0.45 | 5.86 | 3.51 | 0.55 | 8.16 | 5.68 | 0.06 | 13.86 | 11.37 | 0.86 |
| | Infected | 10.93 | 8.37 | | 9.56 | 7.05 | | 6.07 | 4.49 | | 5.80 | 3.34 | | 7.75 | 5.00 | | 13.48 | 11.20 | |
| probing time (s) | Uninfected | 7.61 | 5.78 | < 0.001** | 6.35 | 4.72 | < 0.05* | 5.81 | 4.42 | 0.90 | 6.89 | 4.17 | 0.06 | 6.81 | 5.91 | 0.86 | 14.02 | 14.15 | 0.05 |
| | Infected | 6.38 | 5.11 | | 7.28 | 5.51 | | 5.90 | 4.03 | | 7.06 | 4.35 | | 6.48 | 4.17 | | 10.36 | 9.49 | |
| Average probing bouts per landing | Uninfected | 0.17 | 0.41 | 0.93 | 0.15 | 0.39 | < 0.05* | 0.40 | 0.50 | 0.19 | 0.73 | 0.46 | 0.62 | 0.22 | 0.44 | 0.80 | 0.81 | 1.17 | < 0.05* |
| | Infected | 0.17 | 0.41 | | 0.19 | 0.41 | | 0.37 | 0.50 | | 0.73 | 0.45 | | 0.21 | 0.43 | | 0.93 | 1.07 | |
| total walking distance (mm) | Uninfected | 43.76 | 42.79 | < 0.001** | 38.32 | 34.68 | < 0.05* | 15.17 | 13.40 | 0.30 | 19.69 | 16.21 | 0.14 | 24.64 | 25.21 | 0.45 | 11.00 | 23.66 | 0.10 |
| | Infected | 40.85 | 40.09 | | 36.93 | 34.70 | | 14.93 | 12.77 | | 19.73 | 15.96 | | 23.26 | 22.39 | | 10.88 | 23.15 | |
| mean speed (mm/s) | Uninfected | 3.58 | 1.47 | < 0.05* | 3.64 | 1.57 | < 0.001** | 3.43 | 2.03 | 0.96 | 2.30 | 1.34 | < 0.001** | 3.11 | 1.38 | < 0.001** | 1.13 | 1.99 | < 0.05* |
| | Infected | 3.69 | 1.59 | | 3.51 | 1.56 | | 3.38 | 1.93 | | 2.27 | 1.38 | | 2.99 | 1.35 | | 1.08 | 2.25 | |
| moving | Uninfected | 0.53 | 0.22 | < | 0.54 | 0.23 | < 0.001** | 0.46 | 0.25 | < | 0.28 | 0.19 | < | 0.48 | 0.23 | < | 0.12 | 0.21 | 0.61 |

| | | | | | | | | | | | | | | | | | | |
|---|---|---|---|---|---|---|---|---|---|---|---|---|---|---|---|---|---|---|
| fraction | | | | 0.001** | | | | | | 0.05* | | | 0.001** | | | 0.001** | | |
| | Infected | 0.55 | 0.24 | | 0.51 | 0.23 | | 0.47 | 0.26 | | 0.27 | 0.19 | | 0.45 | 0.22 | | 0.11 | 0.20 |

**Table S5. Epidemiology modelling parameters.**

| Parameter | Full name | Value | | Notes | References |
|---|---|---|---|---|---|
| $S_m$ | Susceptible mosquito population | Malaria | 10000 | Day 0, initial value | - |
| | | Dengue | | | |
| $I_m$ | Infectious mosquito population | Malaria | 0 | Day 0, initial value | - |
| | | Dengue | | | |
| S | Susceptible human population | Malaria | 500 | Day 0, initial value | - |
| | | Dengue | | | |
| I | Infected human population | Malaria | 1 | Day 0, initial value | - |
| | | Dengue | | | |
| R | Recovered human population | Malaria | 0 | Day 0, initial value | - |
| | | Dengue | | | |
| $\mu$ | Mosquito mortality rate | Malaria | 0.095 | - | Anderson & May, 1991 [76] |
| | | Dengue | 0.05 | | Pandey et al. 2013 [77] |
| $B_s$ | Biting rate, susceptible mosquito | Malaria | 0.25 | - | Chitnis et al. 2008 [78] |
| | | Dengue | 0.25 | | Wei Xiang et al. 2022 [32] |
| $B_i$ | Biting rate, infectious mosquito | Malaria | 0.25 | Change subject to the infection-induced behavioral alteration | Chitnis et al. 2008 [78] |
| | | Dengue | 0.25 | | Wei Xiang et al. 2022 [32]. |

| | | | | | |
|---|---|---|---|---|---|
| $E_m$ | Probability, mosquito infection upon exposure to infectious human | Malaria | 0.022 | - | Chitnis et al. 2008 [78] |
| | | Dengue | 0.5 | | Wei Xiang et al. 2022 [32]. |
| $E_h$ | Probability, human infection upon exposure to infectious mosquito | Malaria | 0.36 | - | Chitnis et al. 2008 [78] |
| | | Dengue | 0.5 | | Wei Xiang et al. 2022 [32]. |
| δ | Human recovery rate | Malaria | 180 $days^{-1}$ | - | Filipe et al. 2007 [79] |
| | | Dengue | 5 $days^{-1}$ | | Cummings et al. 2005 [80] |
| ξ | Human immunity loss rate | Malaria | 730 $days^{-1}$ | Approximate value based on previous researches | Doolan et al. 2009 [64]<br>Ghani et al. 2009 |
| | | Dengue | - | | - |

**Table S6. Plasmodium infection rate and average oocyst load per mosquito (oocyst load) of *Anopheles* mosquitoes determined by dissection and qPCR**

| | *Anopheles stephensi - Pf* NF54 | | | *Anopheles stephensi – Pf* NF135 | | | *Anopheles gambiae s.s. – Pf* NF54 | | | *Anopheles coluzzii - Pf* NF54 | | | *Anopheles stephensi - Pv* PvW1 | | |
|---|---|---|---|---|---|---|---|---|---|---|---|---|---|---|---|
| | Dissection & oocyst load | qPCR | Experiment batch | Dissection & oocyst load | qPCR | Experiment batch | Dissection & oocyst load | qPCR | Experiment batch | Dissection & oocyst load | qPCR | Experiment batch | Dissection & oocyst load | qPCR | Experiment batch |
| BuzzWatch experiments | 90%, 7.2 | 80% | Exp 1 | 90%, 9.9 | 95% | Exp 1 | 87.5%, 13.4 | 65% | Exp 1 | 90%, 16.3 | 90% | Exp 1 | 100%, 16.5 | 100% | Exp 1 |
| | 90%, 13.2 | 85% | Exp 2 | 95%, 30.1 | 95% | Exp 2 | 80%, 8.7 | 85% | Exp 2 | 95%, 41.9 | 100% | Exp 2 | 100%, 16.5 | 100% | Exp 2 |
| | 100%, 46.7 | 95% | Exp 3 | 100%, 18.6 | 100% | Exp 3 | | | | | | | | | |
| BiteOscope experiments | 100%, 16.9 | 100% | Exp 1 – Cage 1 | 100%, 31.1 | 100% | Exp 1 – Cage 1 | 100%, 18.4 | 70% | Exp 1 – Cage 1 | 95%, 9.5 | 100% | Exp 1 – Cage 1 | 100%, 16.5 | 100% | Exp 1 – Cage 1 |
| | 100%, 16.9 | 100% | Exp 1 – Cage 2 | 100%, 31.1 | 100% | Exp 1 – Cage 2 | 100%, 18.4 | 70% | Exp 1 – Cage 2 | 95%, 9.5 | 100% | Exp 1 – Cage 2 | 100%, 16.5 | 100% | Exp 1 – Cage 2 |
| | 100%, 16.9 | 90% | Exp 1 – Cage 3 | 100%, 31.1 | 100% | Exp 1 – Cage 3 | 100%, 18.4 | 90% | Exp 1 – Cage 3 | 95%, 9.5 | 90% | Exp 1 – Cage 3 | 100%, 16.5 | 100% | Exp 1 – Cage 3 |
| | 100%, 16.9 | 100% | Exp 1 – Cage 4 | 100%, 31.1 | 100% | Exp 1 – Cage 4 | 100%, 18.4 | 80% | Exp 1 – Cage 4 | 95%, 9.5 | 90% | Exp 1 – Cage 4 | 100%, 16.5 | 100% | Exp 1 – Cage 4 |
| | 100%, 16.9 | 100% | Exp 1 – Cage 5 | 100%, 35.2 | 100% | Exp 2 – Cage 1 | 70%, 10.9 | 80% | Exp 1 – Cage 5 | 90%, 8.9 | 100% | Exp 1 – Cage 5 | 88%, 7.8 | 100% | Exp 1 – Cage 5 |
| | 100%, 9.3 | 100% | Exp 2 – Cage 1 | 100%, 35.2 | 100% | Exp 2 – Cage 2 | 80%, 8.7 | 100% | Exp 2 – Cage 1 | 85%, 6.35 | 90% | Exp 1 – Cage 6 | 100%, 15.2 | 100% | Exp 2 – Cage 1 |
| | 100%, 9.3 | 100% | Exp 2 – Cage 2 | 100%, 35.2 | 100% | Exp 2 – Cage 3 | 80%, 8.7 | 90% | Exp 2 – Cage 2 | 100%, 17.4 | 90% | Exp 2 – Cage 1 | 100%, 15.2 | 90% | Exp 2 – Cage 2 |
| | 100%, 9.3 | 100% | Exp 2 – Cage 3 | 100%, 35.2 | 100% | Exp 2 – Cage 4 | 80%, 8.7 | 90% | Exp 2 – Cage 3 | 100%, 17.4 | 90% | Exp 2 – Cage 2 | 100%, 15.2 | 100% | Exp 2 – Cage 3 |
| | 100%, 9.3 | 80% | Exp 2 – | 100%, 35.2 | 100% | Exp 2 – | 80%, 8.7 | 100% | Exp 2 – | 100%, 17.4 | 100% | Exp 2 – | 100%, 15.2 | 100% | Exp 2 – |

| | | | | | | | | | | | | | | |
|---|---|---|---|---|---|---|---|---|---|---|---|---|---|---|
| | | Cage 4 | | | Cage 5 | | | Cage 4 | | | Cage 3 | | | Cage 4 |
| 100%, 9.3 | 100% | Exp 2 – Cage 5 | 100%, 12.5 | 90% | Exp 3 – Cage 1 | 80%, 8.7 | 70% | Exp 2 – Cage 5 | 100%, 17.4 | 80% | Exp 2 – Cage 4 | 100%, 15.2 | 100% | Exp 2 – Cage 5 |
| 90%, 7.2 | 70% | Exp 3 – Cage 1 | 100%, 12.5 | 90% | Exp 3 – Cage 2 | 80%, 8.7 | 70% | Exp 2 – Cage 6 | 100%, 17.4 | 100% | Exp 2 – Cage 5 | | | |
| 90%, 7.2 | 80% | Exp 3 – Cage 2 | 100%, 12.5 | 100% | Exp 3 – Cage 3 | 80%, 16.9 | 100% | Exp 3 – Cage 1 | 100%, 17.4 | 100% | Exp 2 – Cage 6 | | | |
| 90%, 7.2 | 90% | Exp 3 – Cage 3 | 100%, 12.5 | 100% | Exp 3 – Cage 4 | 80%, 18.4 | 100% | Exp 3 – Cage 2 | 100%, 10.1 | 90% | Exp 3 – Cage 1 | | | |
| 90%, 7.2 | 90% | Exp 3 – Cage 4 | 95%, 19.6 | 90% | Exp 3 – Cage 5 | 80%, 18.4 | 70% | Exp 3 – Cage 3 | 100%, 10.1 | 100% | Exp 3 – Cage 2 | | | |
| | | | | | | 80%, 6.0 | 80% | Exp 3 – Cage 4 | 100%, 10.1 | 100% | Exp 3 – Cage 3 | | | |
| | | | | | | 80%, 6.0 | 90% | Exp 3 – Cage 5 | 100%, 10.1 | 90% | Exp 3 – Cage 4 | | | |
| | | | | | | 80%, 6.0 | 90% | Exp 3 – Cage 6 | 90%, 11.5 | 100% | Exp 3 – Cage 5 | | | |
| | | | | | | 100%, 17.4 | 100% | Exp 4 – Cage 1 | | | | | | |
| | | | | | | 100%, 17.4 | 100% | Exp 4 – Cage 2 | | | | | | |
| | | | | | | 100%, 17.4 | 100% | Exp 4 – Cage 3 | | | | | | |
| | | | | | | 100%, 17.4 | 100% | Exp 4 – Cage 4 | | | | | | |
| | | | | | | 100%, 17.4 | 90% | Exp 4 – Cage 5 | | | | | | |

**Table S7. DENV infection rate of Aedes mosquitoes determined by qPCR**

| | Infection rate | Experiment batch | | Infection rate | Experiment batch |
|---|---|---|---|---|---|
| BuzzWatch experiments | 95% | Exp 1. | BiteOScope experiments | 92% | Exp 1 - cage 1 |
| | 98% | Exp 2. | | 95% | Exp 2 - cage 1 |
| | 92% | Exp 3. | | 94% | Exp 2 - cage 2 |
| | | | | 100% | Exp 3 - cage 1 |
| | | | | 100% | Exp 3 - cage 2 |
| | | | | 100% | Exp 4 - cage 1 |
| | | | | 92% | Exp 4 - cage 2 |
| | | | | 100% | Exp 5 - cage 1 |
| | | | | 96% | Exp 5 - cage 2 |
| | | | | 96% | Exp 6 - cage 1 |
| | | | | 100% | Exp 6 - cage 2 |
| | | | | 83% | Exp 7 - cage 1 |

| | | | | | |
|---|---|---|---|---|---|
| | | | | 96% | Exp 7 - cage 2 |
| | | | | 100% | Exp 8 - cage 1 |
| | | | | 100% | Exp 8 - cage 2 |
| | | | | 100% | Exp 9 - cage 1 |
| | | | | 100% | Exp 9 - cage 2 |
| | | | | 94% | Exp 10 - cage 1 |
| | | | | 95% | Exp 10 - cage 2 |
| | | | | 95% | Exp 10 - cage 3 |
